# B cell-intrinsic requirement for WNK1 kinase in T cell-dependent antibody responses

**DOI:** 10.1101/2021.09.09.459588

**Authors:** Darryl Hayward, Lesley Vanes, Stefanie Wissmann, Sujana Sivapatham, Harald Hartweger, Joshua Biggs O’May, Leon De Boer, Richard Mitter, Robert Köchl, Jens V. Stein, Victor L. J. Tybulewicz

## Abstract

Migration and adhesion play critical roles in B cells, regulating recirculation between lymphoid organs, migration within lymphoid tissue and interaction with CD4^+^ T cells. However, there is limited knowledge of how B cells integrate chemokine receptor and integrin signaling with B cell activation to generate efficient humoral responses. Here we show that the WNK1 kinase, a regulator of migration and adhesion, is essential in B cells for T-dependent antibody responses. We demonstrate that WNK1 transduces signals from the BCR, CXCR5 and CD40, and using intravital imaging we show that WNK1 regulates migration of naive and activated B cells, and their interactions with T cells. Unexpectedly, we show that WNK1 is required for BCR- and CD40-induced proliferation, acting through the OXSR1 and STK39 kinases, and for efficient B cell-T cell collaboration *in vivo*. Thus, WNK1 is critical for humoral immune responses, by regulating B cell migration, adhesion and T cell-dependent activation.

**Summary:** The WNK1 kinase is essential in B cells for T-dependent antibody responses because it is activated by signaling from BCR, CXCR5 and CD40 and regulates B cell migration, adhesion, T-dependent activation, and differentiation into germinal center B cells and plasma cells.

## Introduction

Migration and adhesion play critical roles in B cell physiology. Naïve B cells migrate between lymphoid organs via the bloodstream, entering lymph nodes by extravasating across the endothelium of high endothelial venules using LFA-1 integrin-mediated adhesion triggered by chemokine signals (Girard et al., 2012). Once inside lymphoid tissue, B cells migrate into and within follicular areas under the influence of chemokines such as CXCL13 (Schulz et al., 2016). This continuous movement of B cells allows the cells to locate their cognate antigen and is thus essential for recruitment of B cells into the adaptive immune response.

Migration and cell-cell interactions are also vital during T-dependent B cell activation and subsequent generation of high affinity antibodies after microbial challenge or vaccination (Cyster and Allen, 2019). Binding of cognate antigen to the B cell antigen receptor (BCR) results in internalization of the antigen-BCR complex, degradation of the antigen and, for protein-based antigens, presentation of resulting peptides on MHC class II molecules (Akkaya et al., 2020). Furthermore, binding of antigen to the BCR induces signals that cause movement of B cells from the follicles towards the border of the T cell zone where they interact with CD4^+^ T cells bearing T cell antigen receptors (TCR) specific for the peptide-MHC complex on B cells (Tangye et al., 2013). Subsequent two-way signaling between the B and T cells causes both cell types to become activated and to divide (Crotty, 2015; Petersone et al., 2018). B cell-T cell couples migrate back into the follicle and differentiate into germinal center B (GCB) cells and T follicular helper (Tfh) cells, thereby establishing germinal centers (Mesin et al., 2016; Victora and Nussenzweig, 2012). Within these structures somatic hypermutation of immunoglobulin genes drives affinity maturation, and GCB cells differentiate into antibody-secreting plasma cells and memory B cells. Thus, migration and adhesion play essential roles in B cells during T-dependent activation and need to be closely integrated with activation signals that jointly result in high affinity antibody responses.

WNK1, a member of the WNK-family of protein kinases, is best characterized in kidney epithelial cells where it regulates salt uptake from the urine (McCormick and Ellison, 2011; Shekarabi et al., 2017; Wilson et al., 2001). WNK1 phosphorylates and activates the related OXSR1 and STK39 kinases, which in turn phosphorylate members of the SLC12A-family of ion co-transporters causing net influx of Na^+^, K^+^ and Cl^-^ (Alessi et al., 2014; Thastrup et al., 2012; Vitari et al., 2005; Vitari et al., 2006). Surprisingly, we discovered that in CD4^+^ T cells, WNK1 is activated by signaling from the TCR and CCR7 and that it is a positive regulator of CCR7-induced migration, via OXSR1, STK39 and SLC12A2, and a negative regulator of LFA-1-mediated adhesion (Köchl et al., 2016).

We hypothesized that WNK1 may regulate migration and adhesion of B cells, and thus play an important role in humoral immune responses. We show that WNK1 is activated by signals from the BCR and CXCR5 and is a positive regulator of chemokine-induced migration and a negative regulator of integrin-mediated adhesion. Unexpectedly, we also show that WNK1 is activated by CD40 signaling and that it transduces BCR and CD40 signals via OXSR1 and STK39 that lead to B cell division. Using intravital imaging we find that WNK1 is required for the migration of naive and activated B cells and their interactions with T cells, and show that B cell-intrinsic loss of WNK1 impairs B cell-T cell collaboration, differentiation into GCB cells, and severely reduces T-dependent antibody responses. Thus, WNK1 is critical for humoral immunity because it regulates B cell migration, adhesion, activation and division.

## Results

### B lineage cells express members of the WNK pathway

To evaluate whether WNK1 may regulate B cell migration and adhesion, we first assessed if members of the WNK1 pathway were expressed in B lineage cells. Using our previously generated RNAseq data from 8 different B cell subsets (Brazão et al., 2016), we found that *Wnk1* is the only WNK-family member expressed in any B cell subset (Figures S1A-D). Furthermore, we found that *Oxsr1* and *Stk39*, encoding the related WNK1-activated OXSR1 and STK39 kinases, are expressed in all B cell subsets, albeit expression of *Oxsr1* is higher than *Stk39* (Figure S1E, S1F). Of the seven genes encoding the SLC12A-family of co-transporters, expression of *Slc12a2*, *Slc12a3*, *Slc12a4*, *Slc12a6* and *Slc12a7* was also detected (Figure S1G-M). Thus, B cells express multiple members of the WNK1 pathway.

### Signaling from BCR and CXCR5 activates WNK1

To study the role of WNK1 in B cells, we used three different strategies. Firstly, since constitutive inactivation of *Wnk1* results in embryonic lethality (Zambrowicz et al., 2003), we bred mice with a loxP-flanked (floxed) allele of *Wnk1* (*Wnk1*^fl^), a tamoxifen-inducible Cre recombinase expressed from the ROSA26 locus (*ROSA26*^CreERT2^, RCE) and either a wild type or constitutively deleted allele of *Wnk1* (*Wnk1*^+^ or *Wnk1*^-^) and used bone marrow from the resulting *Wnk1*^fl/+^RCE or *Wnk1*^fl/-^RCE mice to reconstitute irradiated RAG1-deficient mice (Figure S2A). Treatment of these animals with tamoxifen led to efficient deletion of the floxed *Wnk1* allele in B cells 7 d later, hence generating mice with WNK1-deficient B cells and control WNK1-expressing B cells (Figure S2B).

Secondly, to extend the analysis to the function of WNK1 kinase activity we used a *Wnk1* allele expressing a kinase-inactive WNK1-D368A (*Wnk1*^D368A^) (Köchl et al., 2016), reconstituting RAG1-deficient mice with bone marrow from either *Wnk1*^fl/+^RCE or *Wnk1*^fl/D368A^RCE mice (Figure S2A). Treatment of these chimeras with tamoxifen resulted in generation of B cells expressing only WNK1-D368A or control B cells expressing wild-type WNK1. In both cases, loss of WNK1 or its kinase activity caused a drop in the number of mature B cells in the spleen and lymph nodes (Figures S2C, S2D), suggesting that WNK1 activity may be important for B cell survival. Nonetheless, sufficient numbers of WNK1-deficient or WNK1-D368A expressing B cells remained for functional analysis.

Thirdly, it is possible that loss of WNK1 or its kinase activity may cause changes in B cells during the 7 d following the start of tamoxifen treatment, resulting in phenotypes that are not due to an acute requirement for WNK1. To overcome this limitation, we used WNK463, a highly selective WNK kinase inhibitor (Yamada et al., 2016), which allowed investigation of the effect of acute inhibition of WNK1 kinase activity.

WNK1 transduces signals from both the TCR and CCR7 (Köchl et al., 2016), thus we investigated whether, by analogy, WNK1 is activated downstream of the BCR and CXCR5 in B cells. We used phosphorylation of Ser325 on OXSR1 as a readout, as this residue is a direct WNK1 target (Vitari et al., 2005). We found that stimulation of B cells with anti-IgM or CXCL13, the ligand for CXCR5, resulted in a rapid increase in phosphorylated OXSR1 (pOXSR1), which peaked at 10-20 min in response to BCR stimulation and at 2 min in response to CXCL13, with the latter giving a larger response (Figures 1A-F). This increase in pOXSR1 was eliminated by loss of WNK1 or its kinase activity, or by treatment of wild-type B cells with WNK463, demonstrating that both BCR and CXCR5 signaling activates WNK1. Notably, the loss of WNK1 or its kinase activity reduced levels of pOXSR1 to below those detected in unstimulated B cells. Thus, WNK1 kinase has basal activity in resting B cells which is rapidly increased in response to stimulation through the BCR or CXCR5.

**Figure 1.**
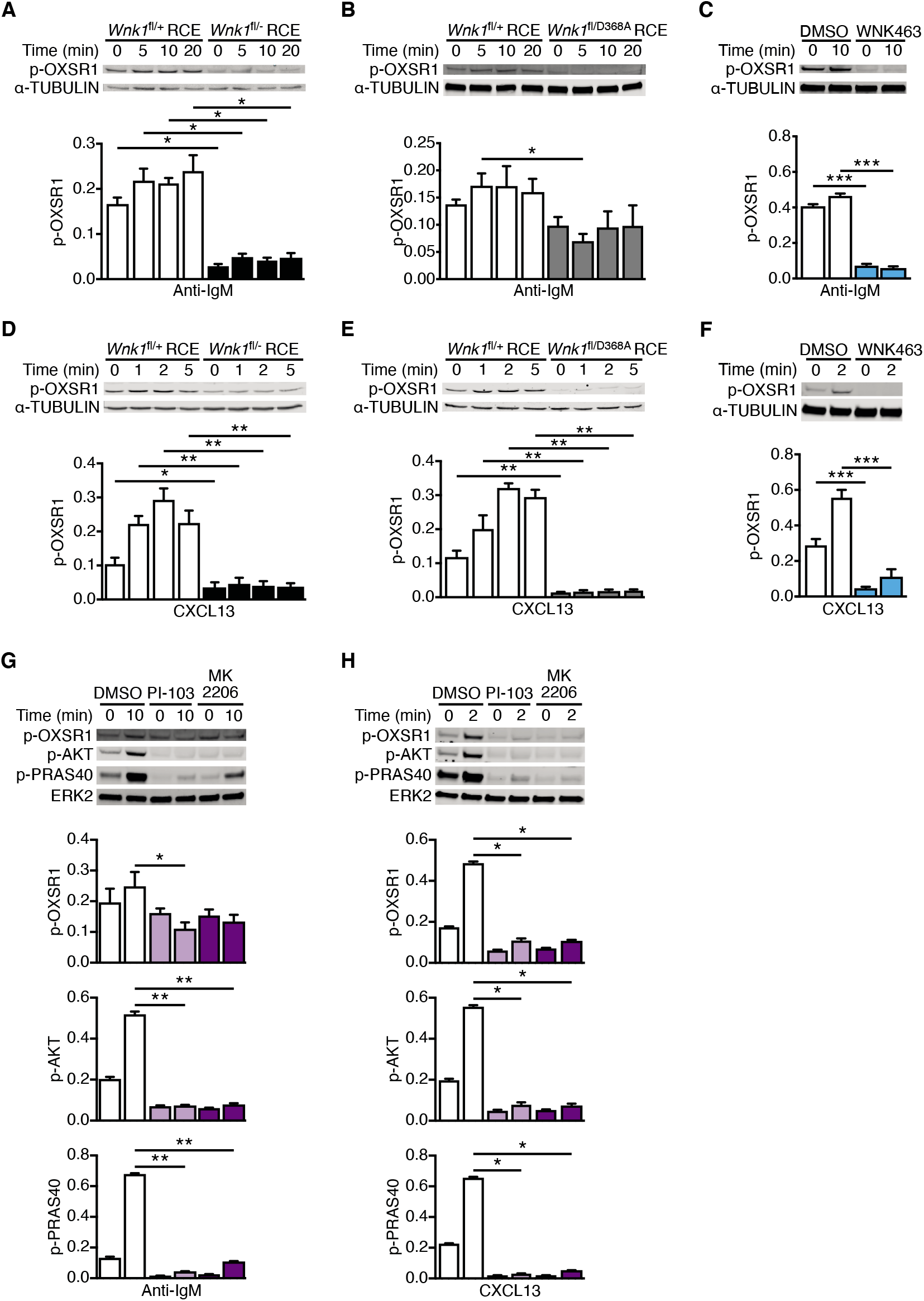
Signaling from the BCR and CXCR5 activates WNK1 via PI3K and AKT. (A-H) Top; immunoblots of cell lysates from mouse B cells stimulated for the indicated times with anti-IgM (A-C, G) or CXCL13 (D-F, H) using WNK1-deficient or control B cells (A, D), B cells expressing kinase-inactive WNK1-D368A or control B cells (B, E), or wild-type B cells treated with vehicle (DMSO), an inhibitor of WNK-family kinases (WNK463) (C, F), a PI3K inhibitor (PI-103) or an AKT inhibitor (MK2206) (G, H), probed with antibodies to phosphorylated OXSR1 (p-OXSR1), α-TUBULIN, phosphorylated AKT (p-AKT), phosphorylated PRAS40 (p-PRAS40) or ERK2. Below; mean±SEM abundance of p-OXSR1, p-AKT and p-PRAS40 in the lanes above, normalized to the α-TUBULIN or ERK2. Mann-Whitney test; * 0.01 < *P* < 0.05, ** 0.001 < *P* < 0.01, *** 0.0001 < *P* < 0.001. Sample sizes: 4 (A, H), 5 (B, D, E, G), 7 (C, F). Data are pooled from two (A, B, D, E, G, H) or three (C, F) independent experiments.

### BCR and CXCR5 signaling activate WNK1 via PI3K and AKT

TCR and CCR7 signaling activates WNK1 via phosphoinositide 3-kinase (PI3K) and AKT (Köchl et al., 2016), thus we investigated if this is also true for BCR and CXCR5 signaling in B cells. Stimulation of B cells with anti-IgM or CXCL13 in the presence of PI-103 and MK2206, inhibitors of PI3K and AKT respectively, resulted in decreased levels of phosphorylated AKT and PRAS40, confirming that the inhibitors were functional (Figures 1G, 1H). Both inhibitors also reduced the BCR- and CXCR5-induced increase in pOXSR1. Thus, both BCR and CXCR5 transduce signals via PI3K and AKT that lead to WNK1 activation.

### WNK1 is a negative regulator of B cell adhesion to ICAM-1

As WNK1 is a negative regulator of LFA-1-mediated adhesion in CD4^+^ T cells (Köchl et al., 2016), we investigated if WNK1 plays a similar role in B cells stimulated via the BCR or CXCR5. We found that in response to stimulation by anti-IgM or CXCL13, B cells deficient in WNK1 or its kinase activity showed substantially higher binding to ICAM-1 and VCAM-1, ligands for the LFA-1 and VLA-4 integrins respectively (Figures 2A, 2B, S2E). Thus, WNK1 is a negative regulator of LFA-1- and VLA-4-mediated adhesion in B cells. Furthermore, loss of WNK1 also induced higher binding to ICAM-1 following treatment of B cells with MnCl_2_ which induces a conformational change in LFA-1 resulting in a high affinity for ICAM-1, demonstrating that WNK1 also regulates this outside-in mode of LFA-1 activation. Analysis of surface proteins showed that WNK1-deficient B cells had normal levels of IgM, slightly increased levels of CD11a, a subunit of LFA-1, and slightly decreased levels of CXCR5 (Figure S2F). The increase in CD11a is much smaller than the increase in adhesion, so it is unlikely to account for the majority of the increased adhesion.

**Figure 2.**
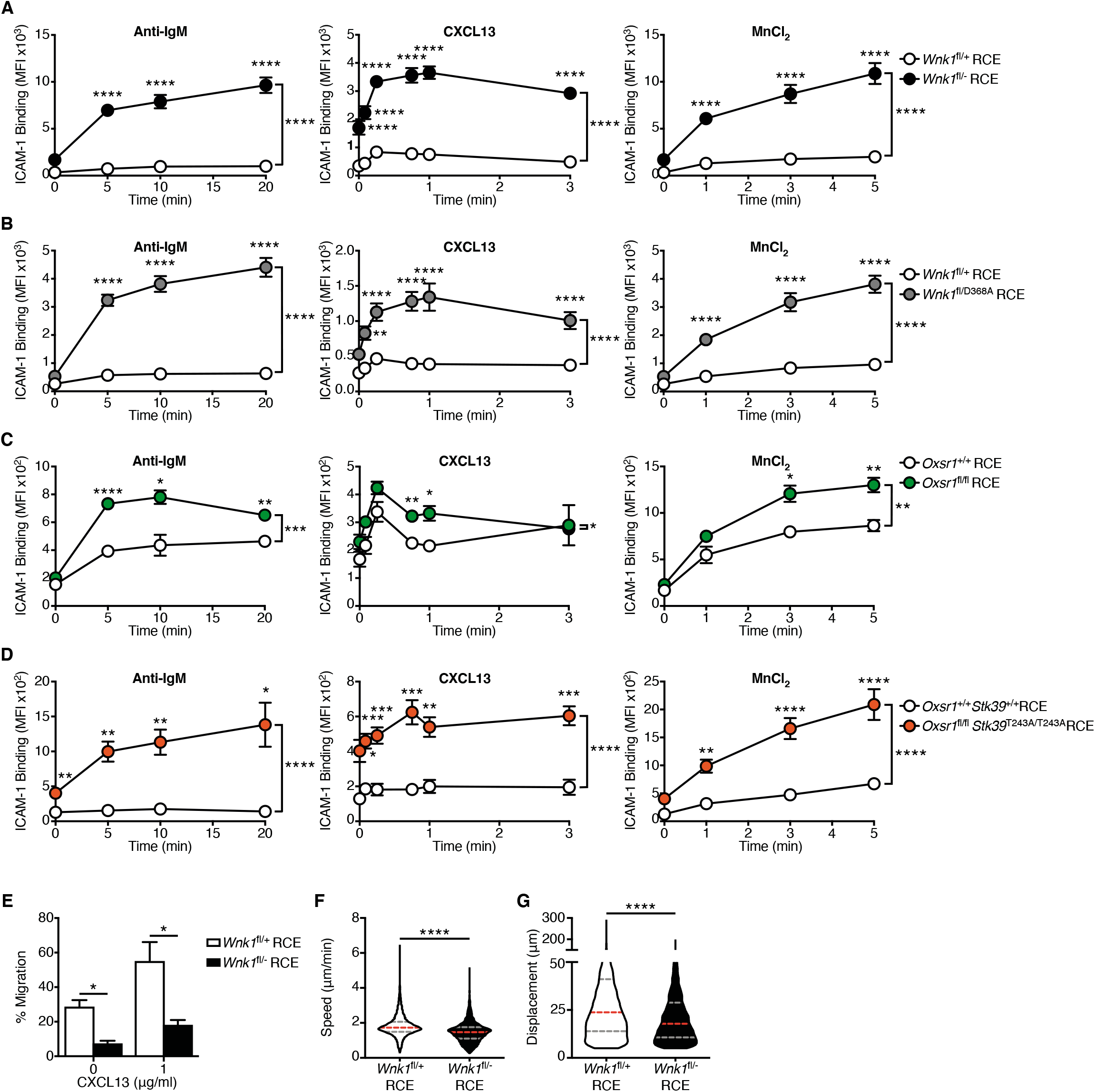
WNK1 regulates B cell adhesion and migration *in vitro*. (A-D) Mean±SEM binding of soluble ICAM-1 complexes to mouse B cells from control B cells and either WNK1-deficient B cells (A), B cells expressing kinase-inactive WNK1-D368A (B), OXSR1-deficient B cells (C) or OXSR1-deficient B cells expressing a non-phosphorylatable mutant of STK39-T243A (D), stimulated with anti-IgM or CXCL13 or treated with MnCl_2_ for the indicated times. (E) Migration of control or WNK1-deficient mouse B cells from the top to the bottom chamber of a Transwell plate in response to CXCL13. (F, G) Violin plots showing mean speed (F) and displacement (G) of control and WNK1-deficient mouse B cells migrating in response to CXCL13. Dashed lines indicate median (red) and 25^th^ and 75^th^ percentiles (grey). Two-Way ANOVA (A-D), Mann-Whitney test (E-G); * 0.01 < *P* < 0.05, ** 0.001 < *P* < 0.01, *** 0.0001 < P < 0.001, **** P < 0.0001. Sample sizes: 11 (A), 6 (B), 5 (C), 9 (D), 4 mutant and 5 control (E), 5245 mutant cells and 7289 control cells (F, G), Data pooled from two independent experiments.

To investigate if OXSR1 or STK39, two well-characterized substrates of WNK1 (Vitari et al., 2005), transduce WNK1 signals that regulate LFA-1 mediated adhesion, we used mice containing floxed alleles of *Oxsr1* (*Oxsr1*^fl^), reconstituting irradiated RAG1-deficient mice with bone marrow from *Oxsr1*^+/+^RCE and *Oxsr1*^fl/fl^RCE mice (Figure S2G). Treatment of these chimeras with tamoxifen resulted in efficient loss of OXSR1, but no change in numbers of mature B cells (Figures S2H, S2I). Analysis of OXSR1-deficient B cells showed that they also had increased ICAM-1 binding in response to treatment with anti-IgM, CXCL13 or MnCl_2_ (Figure 2C). However, the increase was not as large as that seen in the absence of WNK1, suggesting that other WNK1 substrates may be involved, such as STK39. To investigate this potential redundancy between OXSR1 and STK39, we combined the floxed *Oxsr1* allele with an allele of *Stk39* expressing a mutant STK39-T243A protein that cannot be phosphorylated and activated by WNK1 (*Stk39*^T243A^). Reconstitution of irradiated RAG1-deficient mice with bone marrow from *Oxsr1*^+/+^*Stk39*^+/+^RCE and *Oxsr1*^fl/fl^*Stk39*^T243A/T243A^RCE mice (Figure S2G) and subsequent tamoxifen treatment resulted in efficient loss of OXSR1 in the double mutant mice, and again no change in B cell numbers (Figures S2J, S2K).

Analysis of B cells deficient in OXSR1 and expressing STK39-T243A showed that they also had increased binding to ICAM-1 in response to stimulation with anti-IgM, CXCL13 or MnCl_2_, and that the increase was larger than that seen following loss of OXSR1 alone (Figure 2D). These results are consistent with the hypothesis that WNK1 transduces signals from the BCR and CXCR5 via OXSR1 and STK39, leading to negative regulation of LFA-1-mediated adhesion.

### WNK1 is a positive regulator of CXCL13-induced B cell migration *in vitro*

WNK1 regulates migration in different cell types (Köchl et al., 2016; Sun et al., 2006; Zhu et al., 2014). Thus, we hypothesized that WNK1 may also regulate CXCR5-induced migration in B cells. Using a Transwell assay, we found that WNK1-deficient B cells migrated through the Transwell filter much less efficiently in response to CXCL13 (Figure 2E). Furthermore, imaging of CXCL13-induced migration showed that loss of WNK1 resulted in reduced B cell speed and displacement (Figures 2F, 2G). Thus, WNK1 is a positive regulator of CXCL13-induced chemotaxis and chemokinesis.

### WNK1 is required for efficient homing of B cells to lymphoid organs and migration within them

In view of the altered adhesion and migration of WNK1-deficient B cells *in vitro*, and the importance of these processes in B cell trafficking *in vivo*, we examined the ability of the mutant B cells to home efficiently to lymphoid organs. We injected a mixture of control and WNK1-deficient B cells intravenously into mice and 1 h later analyzed their distribution by flow cytometry (Figure 3A). The results showed that fewer WNK1-deficient B cells entered lymph nodes and spleen compared to control B cells, whereas more mutant cells remained in the blood, indicating that WNK1 is required for efficient homing of B cells to lymphoid organs (Figure 3B). Imaging of unsectioned lymph nodes confirmed that fewer mutant B cells had entered the tissue and, in comparison to control B cells, WNK1-deficient B cells were more likely to be located in the lumen of blood vessels or in perivascular areas, and less likely to have entered the parenchyma (Figures 3C, S3A, Video S1). B cells that traffic to the spleen first enter the red pulp and then migrate into the lymphocyte-containing white pulp in response to CXCL13 (Bronte and Pittet, 2013). Histological analysis of the spleen showed that 1 h after transfer, fewer WNK1-deficient B cells were found in both the red and white pulp compared to control B cells, with a relatively larger reduction in the white pulp (Figures 3D, S3B). Taken together these results show that WNK1 is required for B cells to home efficiently to lymphoid organs and to enter the regions of these tissues harboring lymphocytes.

**Figure 3.**
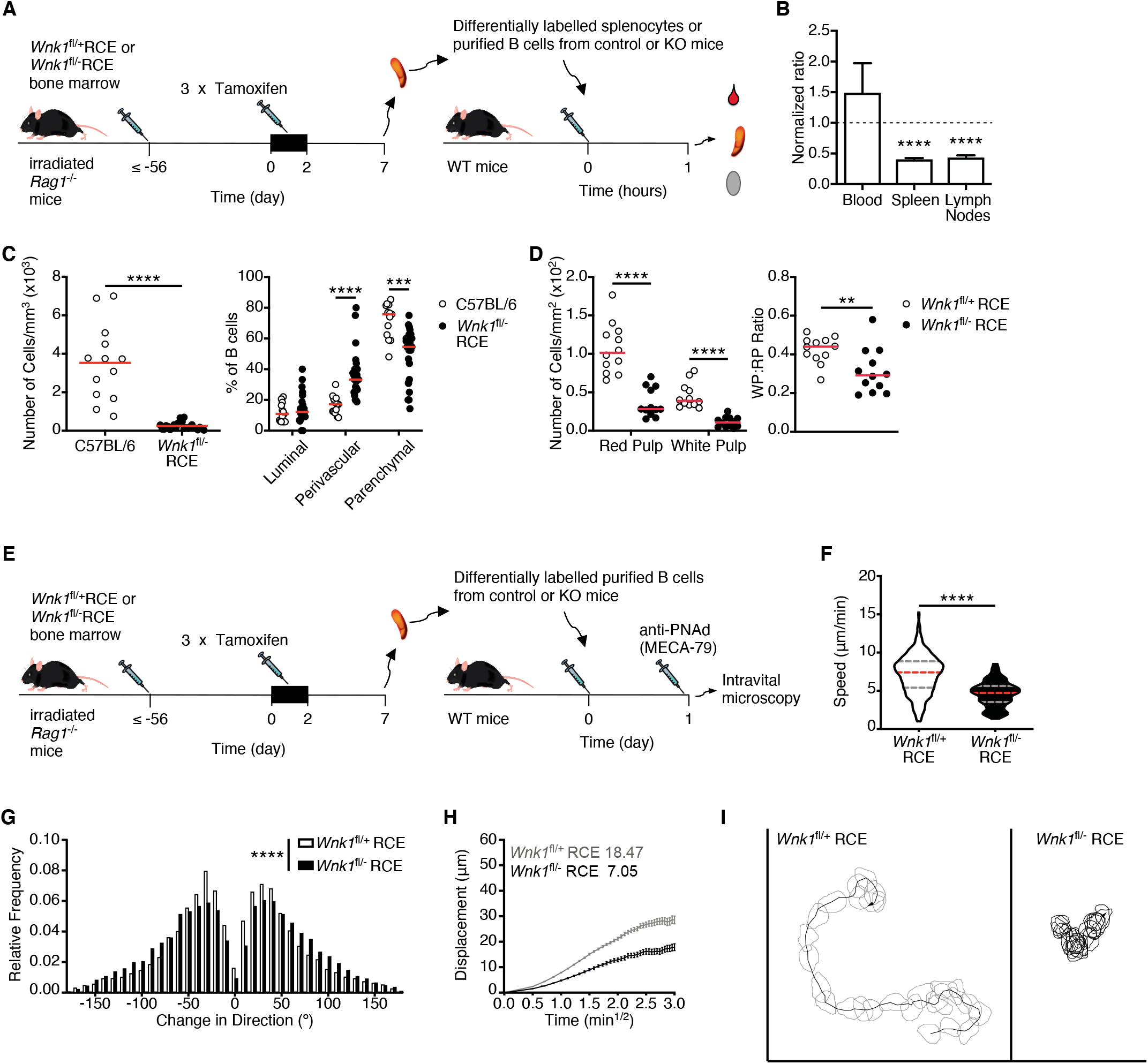
WNK1 is required for efficient homing of B cells to lymphoid tissues, and migration within them. (A) Irradiated RAG1-deficient mice were reconstituted with *Wnk1*^fl/+^RCE or *Wnk1*^fl/fl^RCE bone marrow. At least 56 days later the mice were treated with tamoxifen on 3 consecutive days, splenocytes or splenic B cells were harvested 7 d after start of tamoxifen treatment, control and mutant cells were labelled with two different dyes, mixed at a 1:1 ratio, transferred into wild-type (WT) mice, and 1 h after transfer blood, spleen and lymph nodes were analyzed. (B) Mean±SEM ratio of WNK1-deficient mouse B cells to control B cells in blood, spleen and lymph nodes as described in A. (C) Analysis of 3D histology of lymph nodes showing number of cells per mm^3^ (left), and relative frequency of B cells in the luminal, perivascular and parenchymal regions (right), 40 min after transfer of either WT or WNK1-deficient B cells into WT mice. Each dot is data from one mouse, red line indicates mean. (D) Number of B cells/mm^2^ in the red and white pulp of the spleen (left), and ratio of cells in the white pulp vs red pulp (right), as described in A. (E) Irradiated RAG1-deficient mice were reconstituted with *Wnk1*^fl/+^RCE or *Wnk1*^fl/fl^RCE bone marrow. At least 56 days later the mice were treated with tamoxifen on 3 consecutive days, splenic B cells were harvested 7 d after start of tamoxifen treatment and labelled with two different dyes. Control and WNK1-deficient B cells were transferred into WT mice and 24 h later anti-PNAd (MECA-79) was injected and B cell migration in lymph node follicles was analyzed by intravital microscopy, with results shown in F-I. (F) Violin plot of migration speed; dashed lines indicate median (red) and 25^th^ and 75^th^ percentiles (grey). (G) Relative frequency of change of angle in migration path. (H) Mean±SEM displacement of B cells as a function of the square root of time (time^1/2^); motility coefficients calculated from the slope of the graph are indicated. (I) Typical tracks of a control B cell and a WNK1-deficient B cell, showing cell shapes over 15 min. One-sample t-test (B), Mann-Whitney test (C, D, F, G); ** 0.001 < *P* < 0.01, *** 0.0001 < P < 0.001, **** P < 0.0001. Sample sizes: 8 (B), 13 WT, 26 mutant (C), 12 sections from 3 mice per genotype (D), 5441 WNK1-deficient tracks, 16196 control tracks (F-H). Data pooled from two independent experiments.

Once inside the follicles of lymphoid tissue, B cells continue to migrate in response to CXCL13, a process that allows them to scan cells for presentation of cognate antigen (Akkaya et al., 2020; Cyster and Allen, 2019). To evaluate if WNK1 also regulates this mode of migration, we transferred control or WNK1-deficient B cells into wild-type mice, waited 24 h, and used multi-photon intravital microscopy (MP-IVM) to measure their migration within follicles (Figure 3E). We found that in the absence of WNK1, B cells migrated more slowly, exhibited larger turning angles with reduced track straightness (lower motility coefficient and meandering index) and stopped moving more frequently (increased arrest coefficient) (Figures 3F-3I, S3C, S3D, Video S2). Thus, WNK1 regulates B cell migration within lymphoid tissue.

### WNK1 is required for activation of B cells *in vitro*

Since WNK1 is activated by BCR signaling, it may also contribute to BCR-induced activation, beyond its role in migration and adhesion. To test this possibility, we stimulated control and WNK1-deficient B cells for 3 days with anti-IgM and measured changes in levels of cell surface proteins (activation markers), cytokine secretion, cell division and cell survival. We extended this analysis to include stimulation of the cells through CD40 and TLR4, by treating cells with CD40L and LPS respectively.

In response to BCR stimulation, both control and WNK1-deficient cells upregulated CD69, CD71, CD80, CD86, CD95 and MHC class II (Figure S4A). However, in the absence of WNK1 the upregulation of CD80 and CD95 was higher on day 3 of the culture, whereas upregulation of MHC class II was reduced on days 2 and 3. In response to CD40L stimulation, both control and WNK1-deficient B cells upregulated CD69, CD80, CD86, CD95, ICOSL and MHC class II, but the mutant cells had higher levels of CD69, CD80, CD86 and CD95 on days 1-3, and reduced levels of ICOSL and MHC class II on days 2 and 3 (Figure S4B). Analysis of cytokine secretion showed that in response to anti-IgM WNK1-deficient B cells secreted normal levels of VEGF-A, but higher levels of IL6, IL10 and TNFα, whereas in response to CD40L stimulation secretion of all four cytokines was unaffected (Figure S4C). Thus, in the absence of WNK1, B cells can still be activated by signaling through both the BCR and CD40 but show altered levels of activation markers and cytokines.

In contrast, analysis of cell proliferation showed that WNK1-deficient B cells have greatly reduced cell division in response to either anti-IgM or CD40L stimulation, but normal division in response to LPS (Figures 4A, 4B). WNK1-deficient B cells express unaltered levels of IgM and CD40, so this cannot account for the defects (Figure S2F). We observed reduced recovery of live WNK1-deficient B cells after 3 days of stimulation with anti-IgM, CD40L or LPS compared to control B cells (Figure 4C). To distinguish if this decrease in cell numbers was due to reduced cell division or poorer survival, we calculated numbers of remaining B cells after taking out the effect of cell division. This showed that fewer WNK1-deficient B cells survived in cultures with all three agonists (Figure 4D). Taken together, these results show that WNK1 is required for normal cell division in response to BCR and CD40 but not TLR4 stimulation, and is required for cell survival during B cell activation.

**Figure 4.**
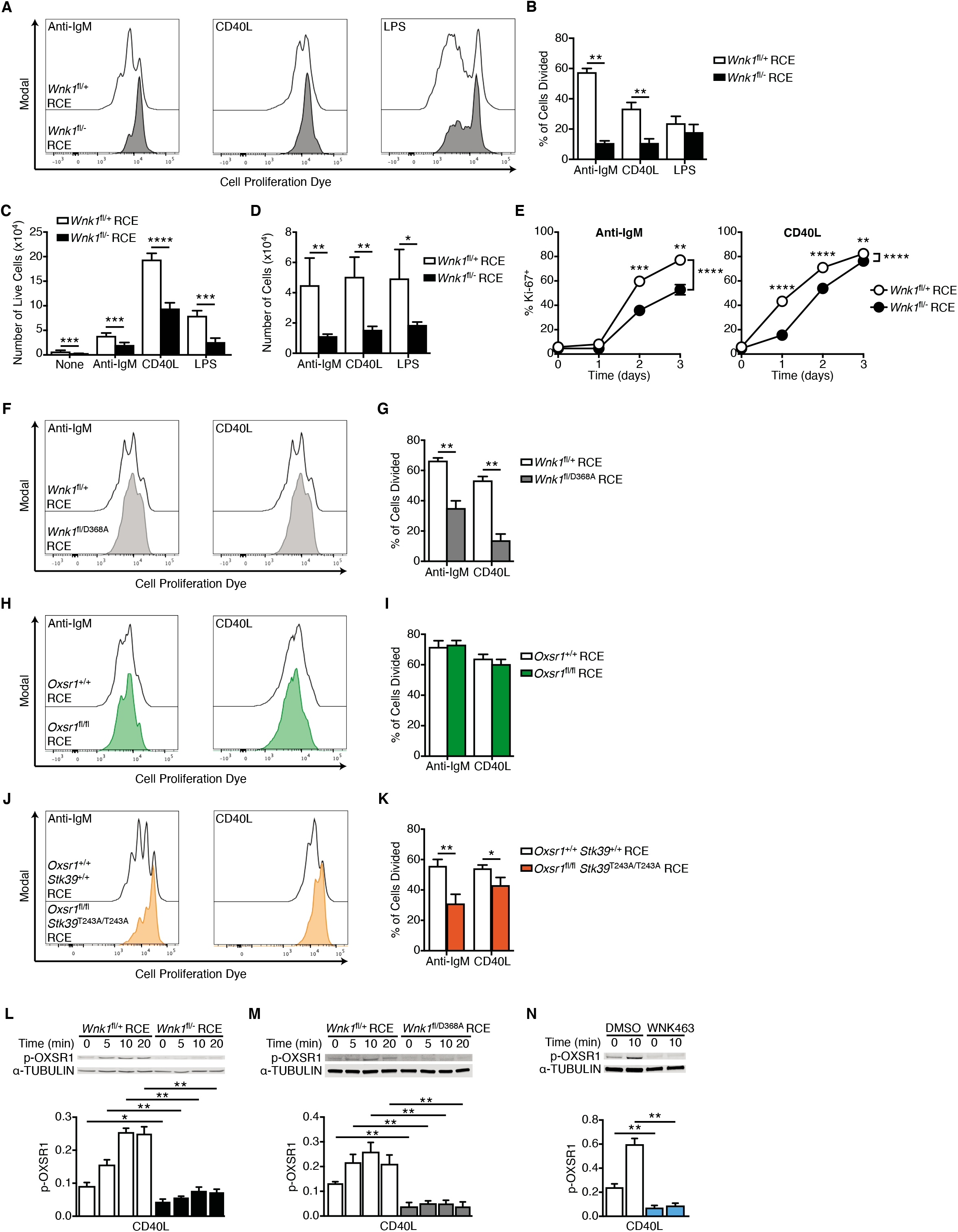
WNK1 is required for B cell activation *in vitro*. (A, B) B cells of the indicated genotypes labelled with Cell Proliferation Dye (CPD) were cultured for 72 h in the presence of anti-IgM, CD40L or LPS. (A) Representative histograms of CPD fluorescence measured by flow cytometry; cell division results in dye dilution. (B) Mean±SEM percentage of B cells that have divided at least once after 72 h stimulation with either anti-IgM, CD40L or LPS. (C) Mean±SEM number of live control or WNK1-deficient B cells after 72 h culture with the indicated stimuli. (D) Mean±SEM number of cells after 72 h culture if there had been no division. (E) Mean±SEM percentage of Ki-67^+^ B cells after stimulation with anti-IgM or CD40L for the indicated times. (F-K) CPD-labelled B cells of the indicated genotypes labelled with CPD were cultured for 72 h in the presence of anti-IgM or CD40L. (F, H, J) Histograms of CPD fluorescence. (G, I, K) Mean±SEM percentage of B cells that have divided at least once after 72 h in response to the indicated stimuli. (L-N) Top; immunoblots of total cell lysates from mouse B cells stimulated for the indicated times with CD40L using WNK1-deficient or control B cells (L), B cells expressing kinase-inactive WNK1-D368A or control B cells (M), or wild-type B cells treated with vehicle (DMSO), or an inhibitor of WNK-family kinases (WNK463) (N), probed with antibodies to p-OXSR1 or α-TUBULIN. Below; graphs of mean±SEM abundance of p-OXSR1 in the lanes above, normalized to α-TUBULIN. Mann-Whitney test (B, C, D, G, I, K-N), two-way ANOVA (E); * 0.01 < *P* < 0.05, ** 0.001 < *P* < 0.01, *** 0.0001 < *P* < 0.001, **** *P* < 0.0001. Sample sizes: 5 WNK1-deficient, 6 control (B, D), 9 WNK1-deficient, 15 control (C), 7 WNK1-deficient, 6 control (E), 4 WNK1-D368A, 6 control (G), 6 (I), 7 (K), 5 (L-N). Data pooled from two (B, D, E, G, I, K-N) or three (C) independent experiments.

We extended this analysis to expression of Ki-67, which is induced when B cells move from the G0 to G1 phase of the cell cycle. Both anti-IgM and CD40L induce Ki-67 expression in B cells with > 80% cells becoming Ki-67^+^ after 3 days of culture (Figure 4E). WNK1-deficient B cells also induce Ki-67 expression, albeit with slower kinetics, and by day 3 of anti-IgM and CD40L stimulation around 50% and 75% of B cells are Ki-67^+^, respectively (Figure 4E). Thus, in the absence of WNK1, B cells are delayed in their entry into G1. In a recent study we showed that WNK1-deficient thymocytes fail to proliferate in response to pre-TCR signals due to defective upregulation of MYC, a transcription factor that induces expression of many proteins required for cell division (Köchl et al., 2020). To investigate if WNK1 plays a similar role in B cells, we measured MYC levels in control and WNK1-deficient B cells in response to anti-IgM and CD40L stimulation. We found that in both cases WNK1-deficient B cells upregulate MYC at least as much as control B cells, suggesting that in B cells WNK1 does not regulate cell division by controlling levels of MYC (Figures S4D, S4E).

To further investigate the mechanism by which WNK1 regulates cell division, we analyzed if WNK1 kinase activity was required for this process. We found that B cells expressing kinase-inactive WNK1-D368A again had greatly reduced cell division in response to anti-IgM and CD40L stimulation (Figures 4F, 4G). Thus, WNK1 kinase activity is required for this activation response. Next, we investigated whether the WNK1 substrates, OXSR1 and STK39 are also involved. Analysis of OXSR1-deficient B cells showed that they divided normally in response to anti-IgM and CD40L (Figures 4H, 4I). In contrast, B cells lacking OXSR1 that also had the STK39-T243A mutation had a reduction in anti-IgM and CD40L-induced cell division, although the defect was not as large as in WNK1-deficient B cells (Figures 4J, 4K). Taken together these results suggest that BCR and CD40 transduce signals via WNK1 to OXSR1 and STK39, which are required for B cells to divide.

### CD40 signaling activates WNK1

Since WNK1 is required for CD40-induced cell division, we investigated if signaling from this receptor activates WNK1. Stimulation of B cells through CD40 resulted in a rapid induction of pOXSR1, peaking around 10 min, which was eliminated by loss of WNK1 or its kinase activity and was also inhibited by WNK463 (Figures 4L-4N). In contrast, treatment of B cells with LPS did not cause any change in pOXSR1 (Figure S4F). Thus, signals from CD40, but not TLR4, induce WNK1 activation. Since both BCR- and CXCR5-induced activation of WNK1 requires PI3K and AKT, we analyzed if the same was true for CD40. Inhibition of PI3K caused no change in CD40-induced pOXSR1, whereas inhibition of AKT resulted in a partial reduction of pOXSR1 (Figure S4G). We conclude that, unlike BCR and CXCR5, CD40 does not activate WNK1 via PI3K, and is only partially dependent on AKT.

### WNK1-deficient B cells activate T cells less efficiently *in vitro*

In view of the defective BCR- and CD40-induced cell division and altered levels of cell surface proteins that are involved in B cell-T cell communication (CD80, CD86, ICOSL and MHC class II), we investigated if WNK1 was required for cognate B cell-T cell interaction and activation. We treated control or WNK1-deficient B cells with beads coated with anti-IgM antibodies and Ovalbumin (OVA), washed out unbound beads and cultured the B cells with OVA-specific OT-II CD4^+^ T cells for 1-3 days. In this assay, crosslinking of the BCR results in the internalization of anti-IgM and OVA-coated beads and the B cells present the OVA 323-339 peptide (OVA_323-339_) on surface MHC class II molecules. These peptide-MHC complexes bind to the TCR on OT-II T cells, triggering the formation of B-T conjugates, and activation and division of the T cells (Figure 5A). In control cultures where the B cells were given no beads (Figure 5B), or beads coated in just anti-IgM or just OVA, or no B cells were present (data not shown), the OT-II T cells did not respond, as measured by upregulation of activation markers (CD25, CD44, CD69) and division. In contrast, in cultures containing WNK1-expressing B cells given beads coated with anti-IgM and OVA, T cells upregulated activation markers and started to divide, demonstrating that T cell activation in this assay requires cognate antigen-driven B cell-T cell interaction (Figure 5B). Notably, in cultures containing WNK1-deficient B cells, the T cells showed lower induction of activation markers, and fewer started to divide. Thus, in the absence of WNK1 in B cells, B-T collaboration is severely impaired.

**Figure 5.**
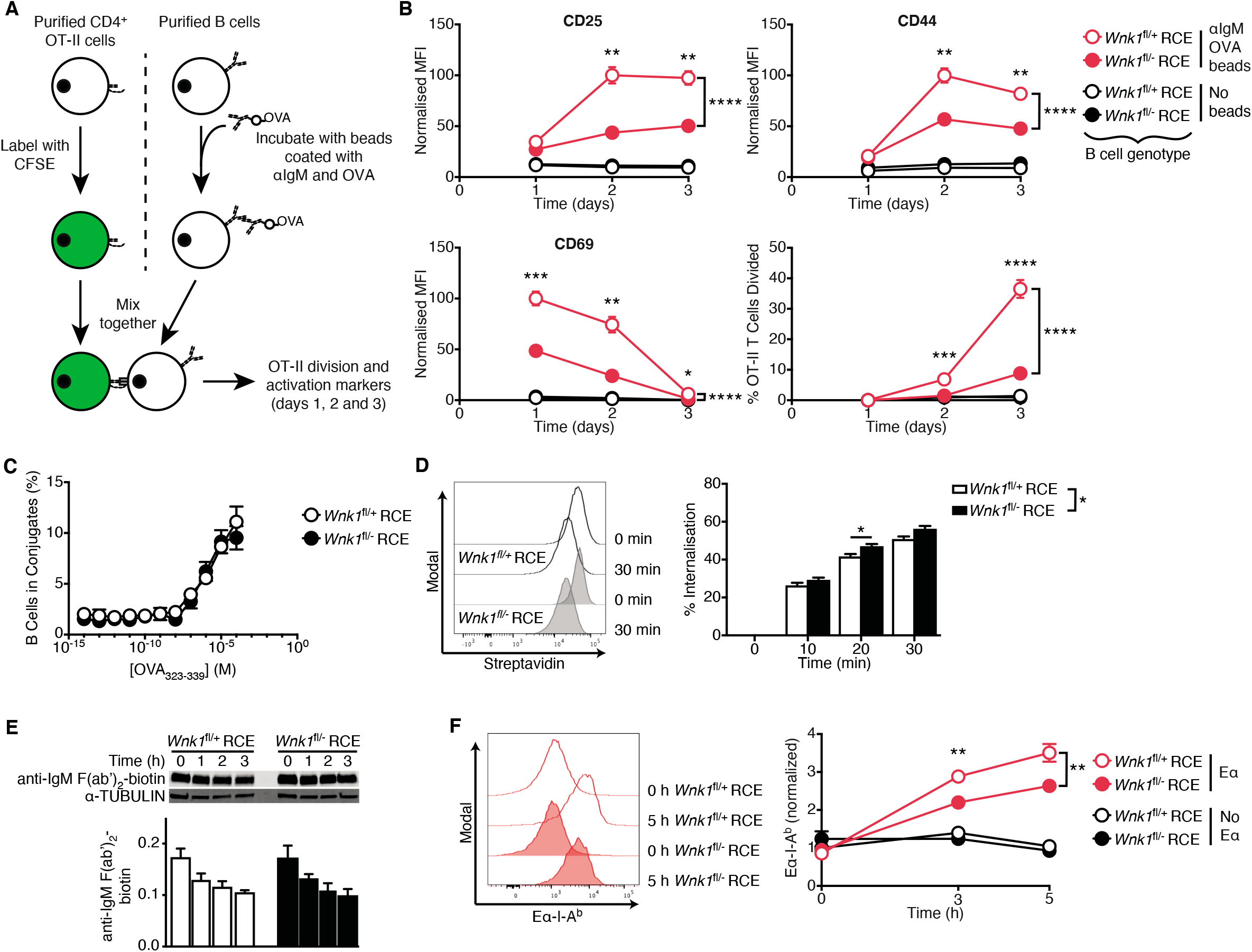
B cells require WNK1 to collaborate efficiently with T cells *in vitro*. (A) Purified CD4^+^ from OT-II mice were labelled with CFSE and cultured with control or WNK1-deficient B cells that have or have not been incubated with beads coated in anti-IgM and OVA for 24, 48 and 72 hours. Activation marker upregulation and cell division was measured using flow cytometry. (B) Mean±SEM normalized MFI of CD25 (top left), CD44 (top right) and CD69 (bottom left), and percentage of cells that have divided at least once (bottom right), of OT-II CD4^+^ T cells cultured for the indicated times with control (open circles) or WNK1-deficient (filled circles) B cells that have been previously incubated with beads coated in anti-IgM and OVA (red) or not incubated with beads (black). MFI was normalized to the maximal response of each individual activation marker (set to 100). (C) Mean±SEM percentage of control or WNK1-deficient B cells that formed conjugates with OT-II T cells as a function of concentration of OVA_323-339_ peptide. (D) B cells of the indicated genotypes were incubated with biotinylated anti-kappa F(ab’)_2_ for the indicated times and residual biotin on the surface revealed with streptavidin as a measure of internalization. Histograms (left) show streptavidin binding, graph (right) shows mean±SEM percentage internalization of the antibody. (E) Immunoblot analysis (top) of total cell lysates from control or WNK1-deficient mouse B cells incubated with biotinylated anti-IgM F(ab’)_2_ for the indicated times, probed with streptavidin to detect biotin, or with an antibody to α-TUBULIN. Graph (bottom) shows mean±SEM abundance of biotinylated anti-IgM F(ab’)_2_ in the lanes above, normalized to α-TUBULIN. (F) Histograms (left) of levels of Eα peptide on I-A^b^ MHC class II on surface of control or WNK1-deficient B cells incubated with Eα peptide-anti-IgM conjugates for the indicated times. Graph (right) shows mean±SEM normalized MFI of Eα-I-A^b^ complex normalized to levels of I-A^b^ MHC class II and to control Eα sample at 0 h (set to 1) as a measure of antigen presentation of control (open circles) or WNK1-deficient (filled circles) B cells incubated with beads coated with anti-IgM and Eα (Eα, red) or just anti-IgM (no Eα, black). Two-way ANOVA (C, D), three-way ANOVA (B, F), Mann-Whitney test (E); * 0.01 < *P* < 0.05, ** 0.001 < *P* < 0.01, *** *P* < 0.001. Sample sizes: 7 WNK1-deficient, 10 control (B), 6 (C, E), 8 (D), 7 (F). Data pooled from 2 independent experiments.

A key event in B-T collaboration is the formation of conjugates in response to presentation of cognate antigen by the B cells to the T cells, a process which is dependent on integrin-mediated adhesion. In view of the role of WNK1 in regulating adhesion, we investigated if conjugation might be perturbed. However, we found no difference in the efficiency of conjugate formation between control and WNK1-deficient B cells (Figure 5C). Once antigen is bound by the BCR, it is internalized by the B cell, degraded, and presented on MHC class II molecules. Analysis of these steps showed that loss of WNK1 did not adversely affect antigen internalization or degradation but caused a partial decrease in the amount of antigen presented on class II molecules (Figure 5D-F). Thus, the defective collaboration of WNK1-deficient B cells with T cells may be due in part to reduced antigen presentation.

### WNK1-deficient B cells fail to mount T-dependent antibody responses

To evaluate the effect of loss of WNK1 on T-dependent antibody responses, we reconstituted irradiated RAG1-deficient mice with a mixture of *Ighm*^μMT/μMT^ bone marrow (80%) which is unable to generate B cells (Kitamura et al., 1991) and either *Wnk1*^fl/+^RCE or *Wnk1*^fl/-^RCE bone marrow (20%). Treatment of the resulting mixed radiation chimeras with tamoxifen results in mice with either WNK1-expressing or WNK1-deficient B cells, whereas most other cells express WNK1. Seven days after the start of tamoxifen treatment, the mice were immunized with NP-conjugated chicken gamma globulin (NP-CGG) in alum, a T-dependent antigen, and blood and splenocytes were analyzed over the following 28 days (Figure 6A). WNK1-deficient B cells were severely impaired in their ability to form NP-specific GCB cells and plasma cells and generated greatly reduced levels of NP-specific IgM and IgG1 in the serum (Figures 6B, 6C, S5A). Thus, WNK1 is required in B cells for T-dependent antibody responses.

**Figure 6.**
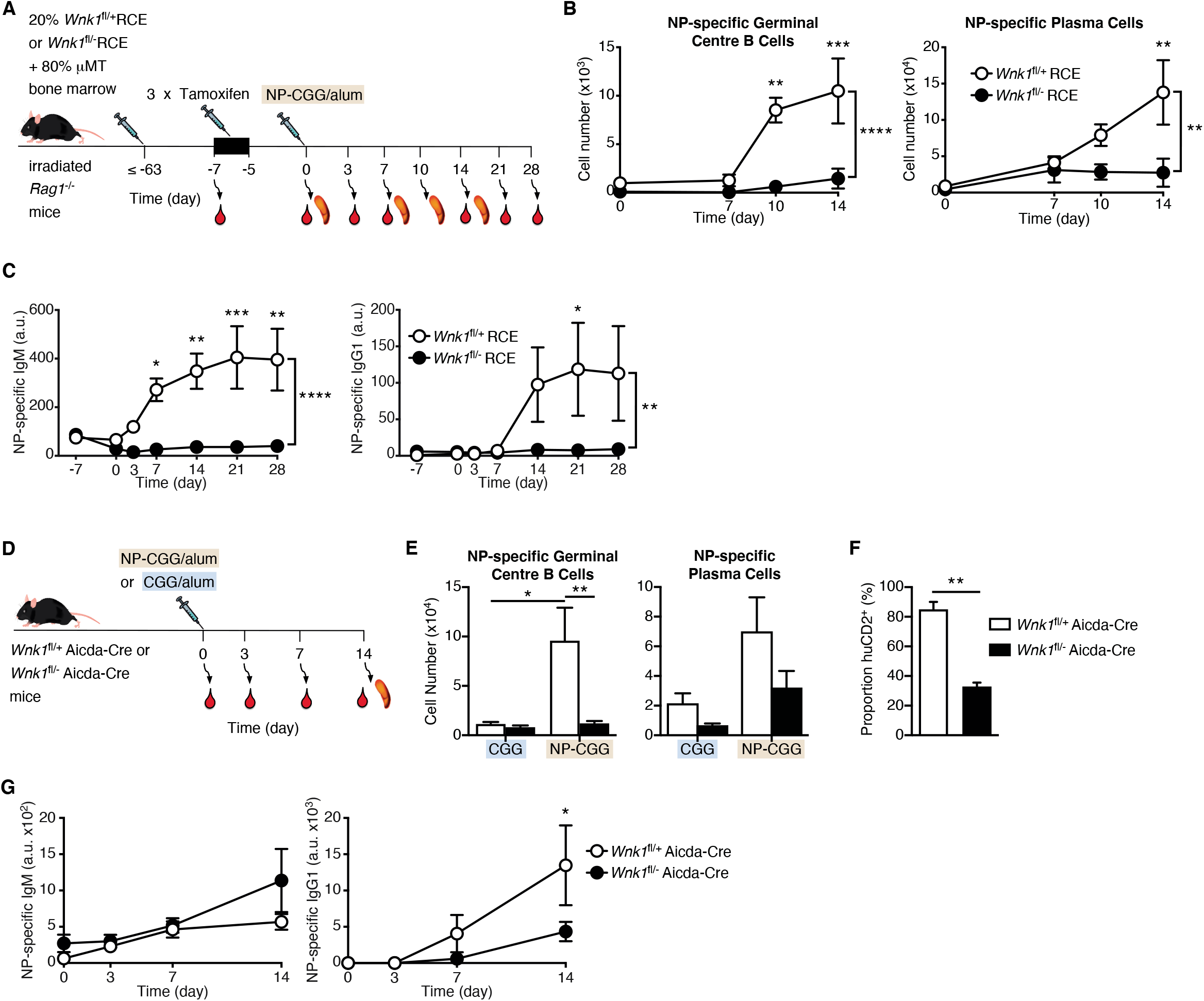
WNK1-deficient B cells fail to mount T-dependent antibody responses. (A) Irradiated RAG1-deficient mice were reconstituted with a mixture of *Wnk1*^fl/+^RCE or *Wnk1*^fl/fl^RCE bone marrow (20%) and μMT marrow (80%). At least 56 days later, blood was taken from the mice and they were treated with tamoxifen on 3 consecutive days, immunized with NP-CGG in alum 7 d after start of tamoxifen treatment and blood and/or spleen were analyzed 0, 3, 7, 10 14, 21 and 28 d later. (B, C) Graphs of mean±SEM numbers of splenic NP-specific germinal center B cells and plasma cells (B) and mean±SEM serum levels of NP-specific IgM and IgG1 (C) in mice treated as described in A. (D) *Wnk1*^fl/+^Aicda-Cre or *Wnk1*^fl/-^Aicda-Cre mice were immunized with NP-CGG or CGG in alum and analyzed 0, 3, 7 and 14 d later. (E-G) Mean±SEM numbers of splenic NP-specific germinal center B cells and plasma cells (E), mean±SEM proportion of germinal center B cells that were hCD2+ (F) and mean±SEM serum levels of NP-specific IgM and IgG1 (G) in mice treated as in D. Two-way ANOVA (B, C, G), Mann-Whitney test (E, F); * 0.01 < *P* < 0.05, ** 0.001 < *P* < 0.01, *** 0.0001 < *P* < 0.001, **** *P* < 0.0001. Sample sizes: 3-4 (B), 4-6 (C), 4-5 (E, F), 5 (G). Data are from 1 (B, C) or 2 independent experiments (E-G).

Since the loss of WNK1 results in lower numbers of B cells (Figure S2C), this could contribute to the reduced antibody response. To evaluate the role of WNK1 without this reduced number of B cells, we used the Aicda-Cre transgene to delete *Wnk1*, since this is only induced in B cells once they are activated. We immunized *Wnk1*^fl/+^Aicda-Cre or *Wnk1*^fl/-^Aicda-Cre mice with CGG or NP-CGG in alum and analyzed the resulting immune response (Figure 6D). We found that loss of WNK1 in activated B cells resulted in a large reduction of NP-specific GCB cells and plasma cells (Figures 6E, S5B). Notably, around 70% of the few remaining GCB cells did not express human CD2, a marker of expression of the Aicda-Cre transgene, implying that these were cells that had most likely not deleted the *Wnk1* gene, further emphasizing the critical role of WNK1 in the differentiation of naïve mature B cells into GCB cells (Figures 6F, S5B). Moreover, in the absence of WNK1 in activated B cells, the levels of NP-specific IgG1 were reduced (Figure 6G), although there was no effect on the levels of NP-specific IgM, most likely because this is generated early in the immune response before the B cells lost WNK1. These results show a cell-intrinsic requirement for WNK1 in B cells for differentiation into GCB cells and plasma cells during a T-dependent immune response.

### B cells require WNK1 to collaborate efficiently with T cells *in vivo*

To understand the role of WNK1 in the early phases of the antibody response, we treated control or WNK1-deficient B cells with beads coated with anti-IgM antibodies, OVA or both and transferred them into mice that had been immunized three days earlier with OVA in alum to provide activated cognate T cells (Figure 7A). Analysis 3 days later showed that B cells treated with beads containing both anti-IgM and OVA responded by upregulating CD69, CD80, CD86, CCR7, ICOSL and MHC class II, as well as inducing expression of Ki-67 and dividing (Figures 7B, 7C, S5C). Little or no response was seen when B cells were given beads coated with just anti-IgM or just OVA, confirming that these responses depend on cognate-antigen driven B cell-T cell interactions. By contrast WNK1-deficient B cells given beads coated with anti-IgM and OVA showed reduced levels of CD69, CD86, ICOSL, MHC class II and Ki-67, and fewer of them had divided. Thus, B cells require WNK1 to upregulate cell surface markers and to divide during B cell activation in response to cognate T cell help *in vivo*.

**Figure 7.**
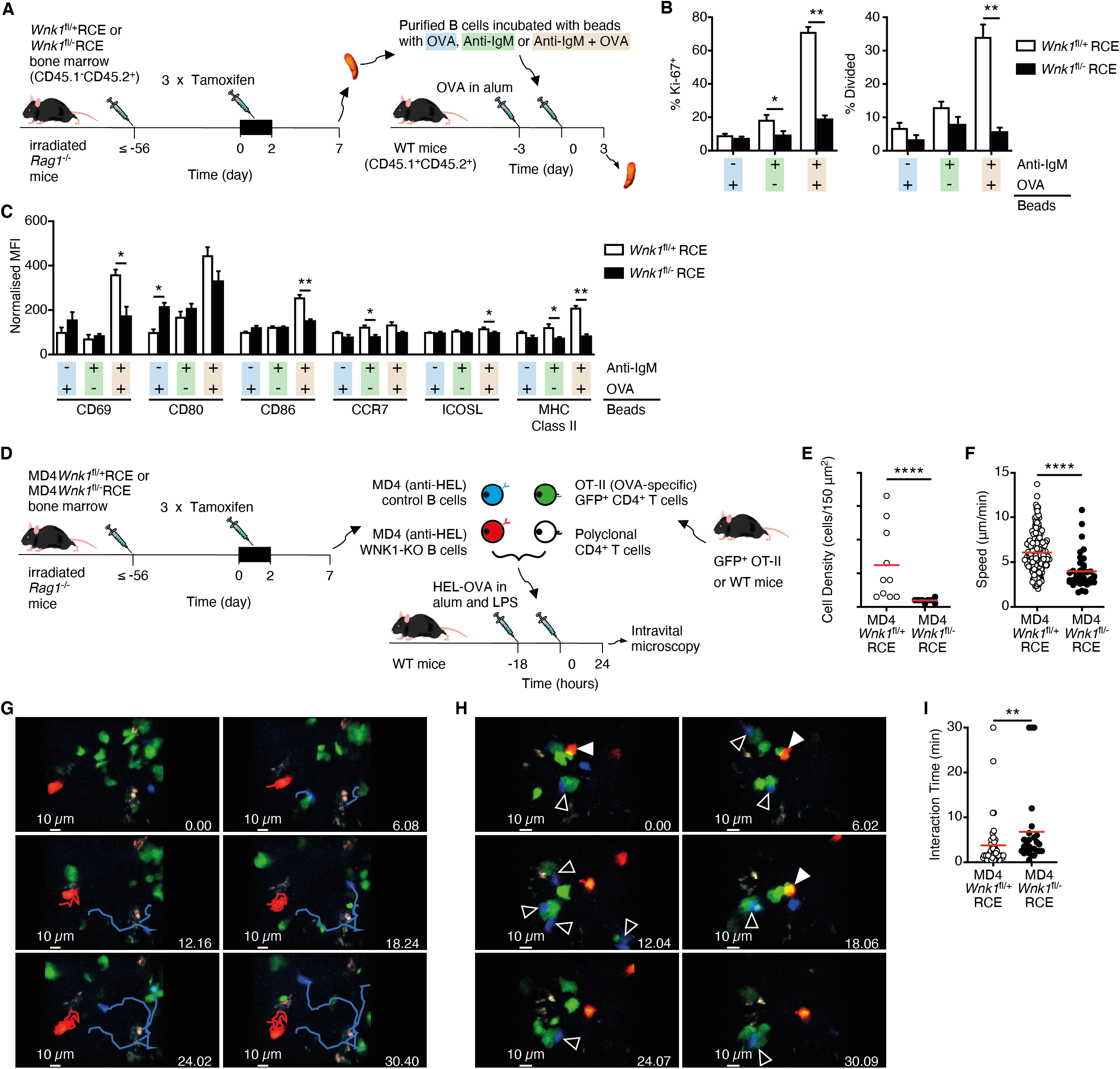
Defective activation of WNK1-deficient B cells by cognate T cells *in vivo*. (A) Irradiated RAG1-deficient mice were reconstituted with *Wnk1*^fl/+^RCE or *Wnk1*^fl/fl^RCE bone marrow (CD45.1^-^CD45.2^+^). At least 56 days later the mice were treated with tamoxifen on 3 consecutive days, splenic B cells harvested 7 d after start of tamoxifen treatment and then labelled with CMFDA and incubated beads coated in anti-IgM and OVA, just anti-IgM or just OVA. The labelled B cells were then transferred into WT mice (CD45.1^+^CD45.2^+^) that were immunized with OVA in alum 3 days prior. 3 days after transfer spleens were harvested and analyzed for activation markers and division by flow cytometry. (B) Mean±SEM percentage of transferred control or WNK1-deficient B cells from mice treated as described in A that express Ki-67 (left) or have divided at least once (right). (C) Mean±SEM median fluorescence intensity (MFI) of the indicated cell surface proteins on transferred control or WNK1-deficient B cells from mice treated as described in A, normalized to the levels on control B cells incubated with beads coated in just OVA (set to 100). (D) Irradiated RAG1-deficient mice were reconstituted with MD4*Wnk1*^fl/+^RCE or MD4*Wnk1*^fl/fl^RCE bone marrow. At least 56 days later the mice were treated with tamoxifen on 3 consecutive days, splenic B cells harvested 7 d after start of tamoxifen treatment, dye-labelled and transferred into C57BL/6J mice that had been immunized in the hock with HEL-OVA in alum and LPS 18 h earlier, along with GFP^+^ OT-II CD4^+^ T cells and dye-labelled polyclonal CD4^+^ T cells. 1 d later the labelled cells were imaged at the B-T border in the draining popliteal lymph nodes by MP-IVM. Results shown in E-I. (E) Density of B cells of the indicated genotype in a 150 μm^2^ field of view. (F) Migration speed; each point represents a single B cell. (G) Time-lapse images showing migration paths of WNK1-expressing (blue) and WNK1-deficient (red) MD4 B cells. (H) Time-lapse images showing interaction of WNK1-expressing (blue, open arrow) or WNK1-deficient (red, filled arrow) MD4 B cells with OT-II T cells (green); polyclonal T cells not shown. (I) Graph of interaction times of MD4 B cells with OT-II T cells. Red lines indicate mean. Mann-Whitney test (B, C, E, F, I); * 0.01 < *P* < 0.05, ** 0.001 < *P* < 0.01, *** 0.0001 < *P* < 0.001, **** *P* < 0.0001. Sample sizes: 7-8 (B, C), 10 fields of view (E), 235 control and 38 mutant cells (F), 46 control and 27 mutant B cells (I). Data are pooled from 2 independent experiments.

These defects in B cell activation could stem from a failure of the B cells to move to the border of the follicle where they can encounter cognate T cells. Alternatively, they may result from impaired interactions with T cells, or defective responses to T cell help. To distinguish these possibilities, we examined the behavior of WNK1-deficient B cells *in vivo* during this early activation phase using MP-IVM. Using B cells expressing the MD4 hen egg lysozyme (HEL)-specific BCR, we transferred dye-labelled WNK1-expressing and WNK1-deficient B cells along with GFP^+^ OVA-specific OT-II CD4^+^ T cells and polyclonal CD4^+^ T cells into mice that had been immunized a day earlier with HEL-OVA in alum in the hock (Figure 7D). One day later we used 4-color MP-IVM to image the transferred lymphocytes at the B cell follicle-T cell zone border in the draining popliteal lymph node. The results showed reduced numbers of WNK1-deficient B cells at the B-T border (Figure 7E), and those that were there migrated more slowly (Video S3, Figure 7F, 7G). These results suggest that activated WNK1-deficient B cells migrate less efficiently to the B-T border. Analysis of B cell-T cell interactions showed that WNK1-expressing MD4 B cells made short duration contacts with OT-II T cells (Video S4, Figures 7H, 7I), typical of cognate B-T interactions (Qi et al., 2008), but not with polyclonal CD4 T cells (not shown). In contrast, the few WNK1-deficient B cells at the B-T border demonstrated altered dynamics, making significantly longer contacts with OT-II T cells, potentially because of their increased integrin-mediated adhesion. Thus, during antigen-driven activation, WNK1-deficient B cells are defective in homing to the B-T border, move more slowly at the border, and make longer interactions with cognate T cells. These changes in the behavior of WNK1-deficient B cells, together with defective CD40-induced responses and reduced antigen presentation are likely to jointly contribute to their profoundly impaired activation, proliferation and subsequent differentiation into GCB cells and plasma cells, and hence their inability to support T-dependent antibody responses.

## Discussion

We have discovered a previously unknown signaling pathway in B cells involving the kinase WNK1. Using a wide variety of *in vitro* and *in vivo* techniques, we found that WNK1 is a critical regulator of multiple aspects of B cell physiology. WNK1 is rapidly activated by signaling from the BCR, CXCR5 and CD40, regulates B cell migration, adhesion, antigen presentation, proliferation and survival, and in its absence from B cells, T-dependent antibody responses are strongly inhibited.

Signaling from the BCR and CXCR5 activates WNK1 via PI3K and AKT, and CD40-induced WNK1 activation is partially AKT-dependent. AKT may directly regulate WNK1, and indeed AKT phosphorylates WNK1 on Thr60 (Jiang et al., 2005; Vitari et al., 2004). However, this phosphorylation does not affect WNK1 *in vitro* kinase activity, or its cellular localization. Thus, it remains unclear how signaling from AKT regulates WNK1 activity, and the identity of other regulators of WNK1 remain unknown.

We found that WNK1 is a positive regulator of CXCR5-induced migration of naïve B cells and that the WNK1-OXSR1-STK39 pathway negatively regulates BCR- and CXCR5-induced activation of adhesion through LFA-1 and VLA-4 integrins. WNK1-deficient T cells have higher levels of active RAP1, a GTPase which is involved in inside-out integrin activation (Alon and Feigelson, 2012). WNK1 may also regulate RAP1 in B cells. It is unknown how the WNK1-OXSR1-STK39 pathway regulates RAP1 activity, but it may act directly or indirectly on RAP1 guanine nucleotide exchange factors or RAP1 GTPase activating proteins.

Unexpectedly, we discovered that WNK1 is required for BCR- and CD40-induced B cell proliferation. This requirement for WNK1 in B cell proliferation is dependent on its kinase activity and is most likely transduced via OXSR1 and STK39 since mutations in these two related kinases resulted in a phenotype similar to WNK1-deficiency. Previous cell line studies reported that WNK1 regulates cell division (Sun et al., 2006; Tu et al., 2011) and our own work in thymocytes showed that WNK1 is required for pre-TCR-induced upregulation of MYC and hence proliferation of thymocytes at the β-selection checkpoint (Köchl et al., 2020). However, the defective proliferation of WNK1-deficient B cells must be due to another mechanism since they upregulate MYC normally.

Some of the best characterized substrates of the WNK-OXSR1-STK39 pathway are the SLC12A-family of ion co-transporters (Shekarabi et al., 2017). Activation of WNK1 leads to phosphorylation and activation of OXSR1 and STK39, which directly phosphorylate multiple SLC12A-family proteins resulting in activation of co-transporters that allow influx of Na^+^, K^+^ and Cl^-^ and inhibition of co-transporters that mediate efflux of K^+^ and Cl^-^ ions. It is possible that the requirement for the WNK1-OXSR1-STK39 pathway in BCR- and CD40-induced B cell activation stems from a need for regulated ion import into the cell, or subsequent water movement, especially as division and migration both require changes, either globally or locally, in cell volume. This is an interesting area for future studies, but these will be challenging given that B cells express five members of the SLC12A-family, which are likely to have redundant functions.

Using an *in vitro* antigen-specific co-culture system, we showed that loss of WNK1 impairs the ability of B cells to activate cognate T cells, potentially due to reduced antigen presentation. While WNK1-deficient B cells internalized and degraded antigen normally, they presented less antigenic peptide on cell surface MHC II molecules. This implies that loss of WNK1 may affect the loading of antigenic peptides onto MHC II molecules in the late endosomal-lysosomal antigen-processing compartment, or subsequent trafficking of peptide-MHC II complexes to the surface (Roche and Furuta, 2015). Several studies have shown that WNK1 regulates vesicle exocytosis and trafficking of membrane proteins to the cell surface (Henriques et al., 2020; Kim et al., 2018; Lee et al., 2004; Mendes et al., 2010; Oh et al., 2007). It will be interesting to investigate if WNK1 regulates exocytosis of peptide-loaded MHC II molecules.

Strikingly, we have shown a B cell-intrinsic requirement for WNK1 in T-dependent antibody responses. WNK1 is required for B cells to receive T cell help and differentiate into GCB cells and plasma cells, because WNK1 regulates B cell migration to the B-T border and communication between B cells and CD4^+^ T cells. WNK1 may also be a critical regulator of GCB cell function as migration and cell-cell interactions are required for GCB differentiation to memory B and plasma cells, both of which provide protection during reinfection. It will be important to further explore how WNK1 regulates humoral responses as this may reveal novel pathways that could be manipulated to improve vaccine responses or suppress generation of pathogenic autoantibodies.

## Materials and Methods

### Mice

Mice with a conditional allele of *Wnk1* containing loxP sites flanking exon 2 (*Wnk1*^tm1Clhu^, *Wnk1*^fl^), with a deletion of exon 2 of *Wnk1* (*Wnk1*^tm1.1Clhu^, *Wnk1*^-^), with a kinase inactive allele of *Wnk1* (*Wnk1*^tm1.1Tyb^, *Wnk1*^D368A^), with a conditional allele of *Oxsr1* containing loxP sites flanking exons 9 and 10 (*Oxsr1*^tm1.1Ssy^, *Oxsr1*^fl^), expressing STK39-T243A (*Stk39*^tm1.1Arte^, *Stk39*^T243A^), with a tamoxifen-inducible Cre in the ROSA26 locus (*Gt(ROSA)26Sor*^tm1(cre/ESR1)Thl^, *ROSA26*^CreERT2^, RCE), with a transgenic Cre recombinase under the control of the *Aicda* promoter followed by an IRES sequence and a truncated human CD2 sequence (Tg(Aicda-cre)9Mbu, Aicda-Cre), with a transgene encoding a T cell receptor that recognizes chicken ovalbumin residues 323-339 in the context of I-A^b^ (Tg(TcraTcrb)425Cbn, OT-II), with a transgene expressing GFP under the control of the human ubiquitin C promoter (Tg(UBC-GFP)30Scha, UBC-GFP), mice deficient for the transmembrane region of the heavy chain of IgM (*Ighm*^tm1Cgn^, *Ighm*^μMT^), or RAG1 (*Rag1*^tm1Mom^, *Rag1*^-^) have been described before (Barnden et al., 1998; de Luca et al., 2005; Kitamura et al., 1991; Köchl et al., 2016; Kwon et al., 2008; Lin et al., 2011; Mombaerts et al., 1992; Rafiqi et al., 2010; Schaefer et al., 2001; Xie et al., 2009). OT-II and UBC-GFP mice were intercrossed to generate GFP^+^ OT-II mice. All strains were bred on a C57BL/6J background. These and (B6 x B6.SJL)F1 mice were maintained in specific pathogen-free conditions at the MRC National Institute for Medical Research and then at the Francis Crick Institute. Both sexes of mice were used, but in each experiment, control and experimental animals were always matched in sex. All animal experiments were carried out under the authority of a Project Licence granted by the UK Home Office and approved by the Animal Welfare Ethical Review Body of the Francis Crick Institute (UK), or animal studies were approved by the Cantonal Committees for Animal Experimentation and conducted according to federal guidelines (Switzerland).

### Radiation chimeras

To generate radiation chimeras, bone marrow cells were harvested from *Wnk1*^fl/+^RCE, *Wnk1*^fl/-^RCE, *Wnk1*^fl/D368A^RCE, MD4:*Wnk1*^fl/+^RCE, MD4:*Wnk1*^fl/-^RCE, *Oxsr1*^+/+^RCE, *Oxsr1*^fl/fl^RCE, *Oxsr1*^fl/fl^*Stk39*^T243A/T243A^RCE, *Ighm*^μMT/μMT^ or OT-II mice, or fetal livers were harvested from E14.5 embryos generated by intercrossing or *Slc12a2*^+/-^ mice. RAG1-deficient animals (5-8 weeks of age) were irradiated with 5Gy using a ^137^Cs-source, and then reconstituted intravenously with at least 1×10^6^ bone marrow cells/recipient. The sex of the donor and recipient mice was always matched. All chimeric animals received Baytril (0.02%, Bayer Healthcare) in their drinking water for at least 4 weeks post-transplantation. If required, 8-20 weeks after reconstitution chimeric mice were injected intraperitoneally with 2 mg/day of tamoxifen (20 mg/ml in corn oil, Sigma) for 3 successive days and analyzed either 7 d after start of tamoxifen treatment for mice containing a *Wnk1*^fl^ allele or treated for 5 successive days and analyzed 21 d after start of tamoxifen treatment for mice containing a *Oxsr1*^fl^ allele.

### Analysis of RNAseq data

Gene expression levels in transcripts per million (TPM) were calculated from previously acquired RNAseq data for B cell subsets (Gene Expression Omnibus, GSE72019) (Brazão et al., 2016) via the RSEM v1.2.31 software (Li and Dewey, 2011), employing STAR v2.5.1b (Dobin et al., 2013) to align reads against the mouse GRCm38 genome assembly with Ensembl release 86 transcript annotations.

### Flow Cytometry

Flow cytometry was carried out using standard techniques with pre-titered antibodies. Antibodies for flow cytometry and cell isolation against the following proteins (all mouse unless otherwise indicated) were obtained from BioLegend, eBioscience, Invitrogen, Jackson Immunoresearch or BD Biosciences (clone names and dilutions indicated in parentheses): B220 (RA3-6B2, 1:200), mouse CD2 (RM2-5, 1:400), human CD2 (RPA-2.10, 1:200), CD4 (RM4-5 or GKL-5, 1:200), CD8α (53-6.7, 1:200), CD11a (M17/4, 1:200) CD11b (M1/70, 1:200), CD11c (N418, 1:200), CD19 (1D3 or 6D5, 1:200), CD23 (B3B4, 1:400), CD25 (PC61.5, 1:400) CD40 (1C10, 1:200), CD43 (eBioR2/60, 1:200), CD44 (IM7, 1:200), CD45.1 (A20, 1:200), CD45.2 (104, 1:200), CD69 (H1.2F3, 1:200), CD71 (R17217, 1:200), CD80 (16-10A1, 1:200), CD86 (GL1, 1:200), CD93 (AA4.1, 1:100), CD95 (Jo2, 1:200), CD138 (281-2, 1:200-300), CXCR5 (L138D7, 1:200), Eα_52-68_ peptide bound to I-A^b^ (eBioY-Ae, 1:200), ICOSL (HK5.3, 1:200), IgD (11-26C, 1:200-400), IgG1 (X56, 1:200-300), IgG2b (polyclonal, 1:500), IgM (RMM-1, 1:200), IgM (goat polyclonal F(ab’)2, 1:300), Ki-67 (SolA15, 1:100), Ly-6G (RB6-8C5, 1:200), Ly77 (GL7, 1:100), MHC class II I-A/I-E (M5/114.15.12, 1:200), TCRβ (H57-597, 1:200). Further reagents: LIVE/DEAD NearIR (1:500), CellTrace^TM^ CFSE, CellTrace^TM^ Violet, CellTracker^TM^ green CMFDA, Cell Proliferation Dye eFluor^TM^ 450, CellTracker^TM^ orange CMTMR, CellTracker^TM^ blue CMAC all from Thermo Fisher Scientific, Inc. and Zombie Aqua (1:500) from Biolegend. For the analysis of Ki-67 expression and of antigen-specific plasma cells, intracellular staining was carried out by incubating single-cell suspensions with antibodies against surface markers, fixing for 20 min with Fix/Perm buffer (BD Biosciences), washing twice with Perm/Wash buffer (BD Biosciences) and incubating with anti-Ki-67 or 4-hydroxy-3-nitrophenylacetyl conjugated to phycoerythrin (NP-PE) (Biosearch Technologies) in Perm/Wash buffer for 30 min. After 2 further washes with Perm/Wash buffer cells were analyzed by flow cytometry.

### Enrichment of splenic B cells

Single-cell suspensions of splenocytes were incubated with biotin-conjugated antibodies against CD11b, CD11c, CD43 and Ly-6G, washed, incubated with Streptavidin-conjugated magnetic beads (Dynabeads, Life Technologies) and cells bound to the beads removed according to the manufacturer’s instructions.

### Immunoblotting

Splenic B cells were rested for 3 h (anti-IgM and CXCL13 stimulations), or not rested (CD40L and LPS stimulations) in IMDM, 5% FCS at 37°C. When indicated, cells were pre-incubated for 1 h at 37°C with either WNK463 (5 µM, HY-100626, Insight Biotechnology), PI-103 (1 µM, Biovision) or MK2206 (2 µM, Cambridge Bioscience) all diluted 1000-fold from stock solutions in DMSO, or with vehicle only (DMSO, 1:1000). Where indicated, the cells were stimulated with either 1 µg/ml recombinant murine CXCL13 (CXCL13, R&D Systems; Biotechne), 10 µg/ml AffiniPure F(ab’)_2_ fragment goat anti-mouse IgM (anti-IgM, Jackson ImmunoResearch Laboratories, Inc.), 10 µg/ml biotin-SP AffiniPure F(ab’)_2_ fragment goat anti-mouse IgM (biotinylated anti-IgM, Jackson ImmunoResearch Laboratories, Inc.), 1 µg/ml recombinant murine CD40L (CD40L, R&D Systems; Biotechne) or 10 µg/ml LPS from *Salmonella minnesota* R595 (LPS, Enzo Life Sciences, Inc.). Subsequent immunoblotting analysis was performed as described previously (Degasperi et al., 2014; Reynolds et al., 2002). The following antibodies and reagents were used for detection of proteins by immunoblotting: anti-pS325-OXSR1/pS383-STK39 (MRC-PPU), anti-OXSR1 (MRC-PPU) anti-α-TUBULIN (TAT-1, Cell Services STP, The Francis Crick Institute), anti-pS473-AKT (193H12, Cell Signaling Technology), anti-pT246-PRAS40 (C77D7, Cell Signaling Technology), anti-ERK2 (D-2, Santa Cruz Biotechnology), anti-MYC (Y69, Abcam). Binding of primary antibodies was detected using IRDye^®^ 800CW-conjugated anti-mouse IgG (LI-COR Biosciences), Alexa Fluor 680-conjugated anti-rabbit IgG (Thermo Fisher Scientific), or Alexa Fluor 680-conjugated anti-sheep IgG (Thermo Fisher Scientific). Biotin was detected using streptavidin-Alexa Fluor 680 (Thermo Fisher Scientific). Fluorescence from the secondary reagents was detected using an Odyssey (LI-COR Biosciences).

### Q-PCR for *Wnk1* mRNA

5×10^5^ splenic B cells from control or WNK1-deficient animals were isolated as described above, total RNA was extracted with an RNAEasy Plus Micro Kit (Qiagen) and cDNA was synthesized with a SuperScript^TM^VILO^TM^ cDNA Synthesis kit (Thermo Fisher Scientific). Samples were analyzed on a QuantStudio3 Real-time PCR system (Thermo Fisher Scientific) using a TaqMan gene expression assay spanning exon1 and exon 2 of *Wnk1* (Mm01184006_m1, Thermo Fisher Scientific). Data was normalized to *Hprt* (Mm03024075_m1, Thermo Fisher Scientific) and analyzed using the comparative threshold cycle method.

### Adhesion assays

Binding of ICAM1 complexes to primary mouse splenocytes was analyzed as described (Konstandin et al., 2006). Either soluble ICAM-1-Fc-F(ab’)_2_ or VCAM-1-Fc-F(ab’)_2_ complexes were generated by diluting APC-labelled goat anti-human IgG F(ab’)_2_ fragments (109-135-098, Jackson Immunoresearch) 1:6.25 with either ICAM-1-Fc (200µg/ml final, Biotechne) VCAM-1-Fc (200µg/ml final, Biotechne) in HBSS and incubated for 30 min in HBSS at 4°C. Splenocytes were rested for 3 h in IMDM, 5% FCS at 37°C, centrifuged and resuspended in HBSS, 0.5% BSA. Each adhesion reaction (10 µl) contained 20×10^6^ cells/ml, 25 µg/ml ICAM-1 or VCAM-1 complex and the appropriate stimulus and was incubated at 37°C for the indicated times. Cells were fixed in PFA for 20 min and binding of ICAM-1 or VCAM-1 complexes to splenic B cells was analyzed by flow cytometry.

### Migration assays

B cell migration was assessed in 96-well Transwell plates, containing polycarbonate filters (5 µm pore size, Corning). Transwell filters were coated overnight with mouse ICAM1-Fc (500 ng/ml in PBS) and blocked with PBS, 2% BSA for over 1 h. The B cells were rested in AB IMDM, 5% FCS for 1 h at 37°C. The receiver plate was filled with RPMI, 0.5% BSA, containing CXCL13 (1 µg/ml) or no chemokine, and 8×10^4^ splenic B cells in RPMI, 0.5% BSA were added to each well of the filter plate. After 3 h at 37°C the filter plate was removed, EDTA was added to each well (40 mM final concentration) and the cells were transferred to 96-well V-bottom plates, spun, resuspended in PBS, 0.5% BSA, and cell numbers determined by flow cytometry. Percentage migration was calculated by dividing the number of cells that migrated through the filter by the total number of cells that had been added to each well.

### Chemokinesis

8-well Chamber Slides (Lab-Tek) were coated overnight with ICAM1-Fc (3 μg/ml) and blocked for 2 h with PBS, 2% BSA. Mouse splenic B cells were labelled with either 1 μM CTV or 1 μM CMFDA in PBS at 37°C, and rested in phenol-red free IMDM, 0.5% BSA, 20 mM HEPES at 37°C for at least 3 h. Cells with different labels were mixed at a 1:1 ratio. 1.5 x 10^5^ cells were added to each chamber and allowed to settle for 20 min in the heat chamber of an Eclipse Ti2 microscope (Nikon, Inc.) set to 37°C. CXCL13 (1 μg/ml) was added to the chamber and three videos (60 frames at 3 frames/min) were generated for each chamber using Eclipse Ti2 and μManager (https://micro-manager.org). Videos were analyzed using the TrackMate (Tinevez et al., 2017) plugin in FIJI (Schindelin et al., 2012). Analysis was limited to cells that appeared in all frames and had a displacement *≥* 5 μm.

### In vivo Homing

C57BL/6J mice were injected intravenously with a 1:1 mixture of splenocytes from *Wnk1*^fl/+^RCE and *Wnk1*^fl/-^RCE bone marrow radiation chimeras, labelled with 1 μM CMFDA or 1 μM CTV. Dyes were swapped between genotypes in repeat experiments. After 1 h, blood, spleen and lymph nodes were harvested, stained with antibodies against B220 and analyzed by flow cytometry to determine the ratio between CMFDA- and CTV-labelled B220^+^ B cells.

### 3D Immunofluorescence

B cells were labeled with CMTMR (5 μM) and transferred i.v. into C57BL/6 recipient mice. After 20 min, further adhesion of B cells to high endothelial venules (HEV) was blocked by i.v. injection of anti-CD62L (Mel-14, 100 µg/mouse; Nanotools) in combination with Alexa Flour 633-conjugated anti-PNAd (MECA-79, 15 μg/mouse, Nanotools) to visualize HEV. After a further 20 min (40 min after B cell transfer), mice were sacrificed and perfused with 10 ml cold PBS and 10 ml cold 4 % PFA. Popliteal, inguinal, axillary and brachial lymph nodes (LNs) were harvested and cleaned from surrounding connective tissues. LNs were fixed overnight in 4% PFA at 4°C. The following day, LNs were embedded in 1.3% low gelling agarose (A-9414, Sigma-Aldrich), dehydrated in 100% methanol (Sigma-Aldrich) overnight, followed by clearing with a benzyl alcohol, benzyl benzoate solution (BABB, Sigma-Aldrich) as described (Boscacci et al., 2010). LNs were imaged by 2-photon microscopy, taking 251 – 501 images from a 300-400 x 300-400 μm field of view (FOV) with z-stacks at 2 μm spacing. Images were visualized with Imaris 9.1.2 software and cells attached to the luminal MECA-79 signal (“luminal”), attached to the abluminal MECA-79 signal (“perivascular”) and cells in the parenchyma of the popliteal, inguinal, axillary and brachial LNs (“parenchymal”) were manually differentiated.

### Histology

C57BL/6J mice were injected intravenously with splenic B cells purified by negative depletion from either *Wnk1*^fl/+^ RCE and *Wnk1*^fl/-^ RCE radiation bone marrow chimeras and labeled with 1 µM CMFDA. After 1 h spleens were harvested and embedded in optimum cutting temperature compound (BDH) and frozen. 8 µm sections were fixed with 4% PFA and blocked with 2% goat serum in PBS. To identify the white pulp sections were stained with anti-MADCAM1 (1:50, MECA-367, eBioscience) and an Alexa Fluor 647-goat anti-rat IgG secondary antibody (1:400, A-21247, Life Technologies). Sections were also counter-stained with DAPI. Images were acquired using an Zeiss Axio Scan.Z1 slide scanner and analyzed by drawing regions of interest around the spleen and the white pulp areas and cells were counted using FIJI. Numbers of cells were first normalized to the red and white pulp area, respectively and a ratio of cell density in the white pulp to red pulp was then calculated.

### Intravital Microscopy of Naïve B cells

Splenic B cells were isolated using a mouse B cell isolation kit (STEMCELL Technologies) according to the manufacturer’s protocol. B cells of each genotype were labelled either with 5 μM CMTMR or 20 μM CMAC for 20 min at 37°C, 5% CO_2_. and mixed at a 1:1 ratio. The mixture was transferred intravenously into sex-matched C57BL/6J mice. 24 h after transfer, the popliteal LN of recipient mice was surgically exposed for two-photon microscopy (2PM) as described (Moalli et al., 2014). Immediately prior to imaging, HEVs were labelled by intravenous injection of Alexa Fluor 633-conjugated anti-PNAd (MECA-79). 2PM imaging was performed by using a Trimscope system equipped with an Olympus BX50WI fluorescence microscope and a 20X (NA 0.95, Olympus) or 25X objective (NA 1.10, Nikon), controlled by ImSpector software (LaVision Biotec). A MaiTai Ti:Sapphire laser (Spectra-Physics) was tuned to 780 nm for excitation of the fluorophores. To perform four-dimensional analysis of cell behavior, a 250 x 250 µm FOV with z-steps of 4 µm spacing was recorded every 20 s for 20 - 30 min. Vivofollow software allowed a continuous drift offset correction in real time using fine pattern matching during 2PM imaging (Vladymyrov et al., 2016). Cell migration analysis was performed using Imaris 9.1.2 software (Bitplane). Median speeds, meandering index and turning angles (defined as the angle between the two velocity vectors before and after a measurement time point) were calculated from the (x,y,z) coordinates of the transferred B cell centroids. The arrest coefficient was calculated as the percentage of time per track that a cell moved < 4 μm/min, using a custom script for Matlab (MathWorks) (Moalli et al., 2014). The motility coefficient was calculated from the slope of a graph of displacement against time^1/2^.

### Activation, Proliferation and Survival Assays

Splenic B cells were purified by negative depletion as described above and resuspended in DMEM, 100 μM non-essential amino acids, 20 mM HEPES buffer (all from Thermo Fisher Scientific), 10% FCS, 100 U/ml Penicillin (Sigma-Aldrich; Merck KGaA), 100 μg/ml Streptomycin (Sigma-Aldrich; Merck KGaA), 2 mM L-Glutamine (Sigma-Aldrich; Merck KGaA) and 100 μM 2-mercaptoethanol (Sigma-Aldrich; Merck KGaA) (DMEM+). To measure proliferation, cells were labelled with Cell Proliferation Dye eF450 (10 μM, Thermo Fisher Scientific) in PBS for 10 min at 37°C before quenching with FCS-containing media. Cells were cultured in flat bottom 48-well plates (Corning, Inc.) at 10^6^ cells/well and stimulated with either 10 μg/ml anti-IgM F(ab’)_2_, 1 μg/ml CD40L, or 10 μg/ml LPS at 37°C for indicated times. After culture the cells were subjected to centrifugation, and the supernatants removed and stored at −80°C for Luminex analysis. The number of live cells, surface protein levels and proliferation were determined using flow cytometry. The number of cells remaining if there had been no divisions (N_0_) was calculated as 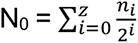 where n_i_ is the number of cells that had undergone *i* divisions, and *z* is the largest number of divisions observed in a given experiment.

### Luminex Cytokine Secretion

A 4-Plex ProcartaPlex kit (Thermo Fisher Scientific, Inc.) was used to measure IL-6, IL-10, TNF-α and VEGF-A using Luminex-based technology according to the manufacturer’s instructions on a Bio-Plex 200 (Bio-Rad Laboratories).

### Antigen Presentation to OT-II T Cells

0.2 μm streptavidin coated polystyrene microspheres (Bangs Laboratories) were incubated at 37°C for 1 h with biotinylated anti-IgM at a final concentration of 0.561 μg/10^9^ microspheres and biotinylated OVA (Nanocs, Inc.) at a final concentration of 0.219 μg biotin-OVA/10^9^ microspheres. Conjugated microspheres were counted using a ZE5 cell analyzer (Bio-Rad Laboratories, Inc.) and resuspended in DMEM+ at a bead concentration of 10-25 x 10^8^/ml. Splenic B cells were isolated as described before and naïve splenic CD4^+^ T cells were isolated from OT-II mice were isolated using biotinylated antibodies against B220, CD8, CD11b, CD11c, CD19, CD25 and CD44. T cells were labelled with 1 μM CFSE. Splenic B cells were incubated with conjugated microspheres at a ratio of 10-25 beads per B cell for 30 min at 37°C. Excess microspheres were washed away. Co-cultures of 2 x 10^5^ B cells and 10^5^ OT-II CD4^+^ T cells were cultured in 96-well U-bottom plates at 37°C for 24, 48 and 72 h. Flow cytometry was used to determine proliferation and upregulation of activation markers.

### B-T Conjugation

Splenic B cells were isolated using negative depletion as described above, and naïve splenic and lymph node CD4^+^ T cells from OT-II bone marrow chimeras were isolated using negative depletion as described above. The B cells were incubated with the indicated concentrations of OVA_323-339_ for 30 min at 37°C in FCS-free medium. The peptide-loaded B cells were mixed with the OT-II T cells at a 1:1 ratio and incubated at 37°C for 1 h, fixed with 4% PFA and pipetted 12 times to remove non-specific conjugates. Flow cytometry was used to determine conjugation frequency defined as the frequency of B cells in a conjugate with a T cell.

### Antigen Internalization

Goat F(ab’)_2_ anti-mouse Igκ (Southern Biotech) was biotinylated with 20-fold molar excess of NHS-LC-LC-biotin (Thermo Fisher Scientific, Inc.) and labelled with Cy3 monoreactive dye pack (GE Healthcare) in sodium carbonate buffer (biotinylated anti-κ-Cy3). Excess dye was removed using Zeba 7K MWCO desalting columns (Thermo Fisher Scientific, Inc.). Splenocytes were incubated with biotinylated anti-κ-Cy3 for 30 min on ice. Samples were incubated at 37°C for the times indicated before fixation with 2% PFA on ice for 20 min. Cells were stained with fluorescently-labelled streptavidin to label the biotinylated anti-κ-Cy3 on the surface. Flow cytometry was used to determine antigen internalization defined as the reduction in the amount of biotinylated anti-κ-Cy3 remaining on the surface as a percentage of the amount on the surface of cells that had been left on ice.

### Eα Peptide Presentation

0.2 μm Dragon Green streptavidin microspheres (Bangs Laboratories, 10 mg/ml) were incubated at 37°C for 1 h with biotinylated anti-IgM at a final concentration of 1.52 μg/10^9^ microspheres, and with or without 3.03 μg biotin-Eα peptide (Biotin-GSGFAKFASFEAQGALANIAVDKA-COOH, Crick Peptide Chemistry)/10^9^ microspheres. Conjugated microspheres were resuspended in DMEM+ at a bead concentration of 40 μg/ml. 1.3 x 10^6^ splenic B cells were incubated with 4 μg conjugated microspheres for 30 min at 37°C. Excess microspheres were washed away, cells were incubated at 37°C for the indicated times and fixed with a final concentration of 2% formaldehyde (Thermo Fisher Scientific, Inc.). Quantities of Eα peptide on the I-A^b^ MHC class II molecule and total I-A^b^ were determined using flow cytometry.

### NP-CGG Immunization

Mice were injected i.p. with 50 μg of either chicken gamma globulin (CGG) or NP-CGG (BioSearch Technologies) with 25% Alum (Thermo Fisher Scientific, Inc.) in PBS. The immune response was assessed by either withdrawing blood from superficial veins of the mouse tail or the mice were sacrificed at the indicated time points and spleens were used for flow cytometric analysis.

### ELISA

Blood harvested from the superficial tail vein was left to clot for at least 30 min before centrifugation at 17000 g for 10 min at room temperature. The supernatant was transferred to a new tube and centrifuged again at 17000 g for 10 min. Finally, the supernatant was transferred to a new tube and stored at −80°C until the ELISA was performed. 96-well Maxisorp Immunoplates (Nalge Nunc International Corporation) were coated with 5 μg/ml NP_20_-BSA (Santa Cruz Biotechnology, Inc.) overnight at 4°C and then washed five times with PBS, 0.01% Tween-20. Plates were blocked with 3% BSA in PBS for 2 h at room temperature and washed twice with PBS, 0.01% Tween-20. Serum was diluted 1:100 in PBS for the IgM ELISA and 1:1000 for the IgG1 ELISA and a further serial dilution was performed with 2-fold dilution steps. Diluted sera were added to the coated plates and incubated overnight at 4°C, followed by three washes with PBS, 0.01% Tween-20. Biotin-conjugated goat anti-mouse IgM (1.6 μg/ml, CALTAG laboratories, Thermo Fisher Scientific, Inc.) or biotin-XX conjugated goat anti-mouse IgG1 (4 μg/ml, Invitrogen; Therrmo Fisher Scientific, Inc.) were added to their respective plates and incubated at room temperature for 2 h. After three washes, plates were incubated with peroxidase labelled streptavidin (1 μg/ml, Vector Laboratories) for 2 h at room temperature. Plates were washed five times before incubation with 1x TMB ELISA substrate solution (eBioscience; Thermo Fisher Scientific, Inc.) and the reaction was stopped with 1 M H_2_SO_4_. Signals were quantified using absorption at 450 nm on a SpectraMax 190 (Molecular Devices LLC). Data was analyzed using Microsoft Excel and GraphPad Prism 8 (GraphPad). Values in the linear portion of the response curve were used to calculate concentrations of NP-specific antibodies in arbitrary units.

### *In Vivo* Activation and Proliferation

(B6 x B6.SJL)F1 mice were injected i.p. with 50 μg of ovalbumin (Invivogen) with 25% Alum (Thermo Fisher Scientific, Inc.) in PBS. Three days later, splenic B cells were isolated by negative depletion, labelled with CMFDA (1 μM, Thermo Scientific, Inc.) and incubated with microspheres conjugated with either OVA, anti-IgM or both as described above. These cells were then injected intravenously into the pre-immunized (B6 x B6.SJL)F1 mice at 500,000 cells per mouse. Mice were sacrificed three days later and spleens were analyzed by flow cytometry.

### HEL-OVA conjugation

HEL and OVA (both from Sigma) was mixed in a 1:1 ratio (w/w) in PBS containing 0.023% glutaraldehyde and stirred at room temperature for 1 h. After removing precipitate by centrifugation, the solution was dialysed against PBS (Dialysis tubes 8 kDa molecular weight cutoff Mini Dialysis kit, Cytiva) overnight at 4°C as described (Allen et al., 2007). HEL-OVA conjugation was confirmed by SDS-PAGE and silver staining. Conjugated HEL-OVA solution was further purified from unconjugated proteins using a 50 kDa Amicon filter tube (Merck).

### Intravital Microscopy of B cell-T cell interactions

Splenic B cells were isolated from bone marrow chimeras (MojoSort™ Mouse Pan B Cell Isolation Kit; Biolegend). Control and WNK1-deficient B cells were labelled with CellTracker blue (CMAC, 20 µM) and CellTracker orange (CMTMR, 10 µM; both Invitrogen) respectively, for 20 min at 37°C. CD4^+^ T cells were isolated from WT C57BL/6J mice and from GFP^+^ OTII mice (EasySep Mouse CD4^+^ T Cell Isolation Kit; Stemcell). WT polyclonal CD4^+^ T cells were labelled with Cell Proliferation Dye eFluor670 (10 µM; eBioscience) for 20 min at 37°C. Control and WNK1-deficient B cells (5×10^6^ each) were transferred i.v. in a 1:1 ratio together with GFP^+^ OT-II TCR transgenic CD4^+^ T cells and polyclonal CD4^+^ T cells (3 x 10^6^ each) into recipient mice that had received 15 µg HEL-OVA, 0.2 µg LPS in Alum (total volume of 20µl) s.c. in the foot hock 18 h earlier. At 24 h post cell transfer, intravital imaging of reactive popliteal lymph nodes was performed on a TrimScope (LaVision Biotec) equipped with a MaiTai NIR Laser (Spectraphysics) tuned to 780 nm and a Nikon 25X objective with NA 1.0 (Ficht et al., 2019). For analysis of cell migration and interactions, z-stacks of 11 images from 150 – 250 µm^2^ areas were recorded every 30 s for 30 min at the B cell follicle border. Image sequences were analyzed by semi-automated tracking using Imaris (Bitplane) and by visual inspection for interaction time.

### Statistical Analysis

All statistical comparisons were carried out using the nonparametric two-tailed Mann-Whitney test or a Two-Way ANOVA test as detailed in Figures (Prism 8, GraphPad).

## Author contributions

DH, LV, SW, SS, HH carried out experiments and analyzed data, DH and VLJT designed the study and wrote the manuscript, RK, JVS and VLJT supervised the work.

## Acknowledgments

We thank Edina Schweighoffer for technical advice. We thank Pavel Tolar, George Kassiotis, Edina Schweighoffer, Daisy Luff and Fábio Ribeiro Rodrigues for critical comments on the manuscript. We thank the Flow Cytometry, Experimental Histopathology, Crick Advanced Light Microscopy, Cell Services, Peptide Chemistry, Media Kitchen and Biological Research Facilities of the Francis Crick Institute for scientific support, reagents and animal husbandry. We thank Chou-Long Huang, Dario Alessi and Sung-Sen Yang for mouse strains. VLJT was supported by the UK Medical Research Council (Programme U117527252) and by the Francis Crick Institute which receives its core funding from Cancer Research UK (FC001194), the UK Medical Research Council (FC001194), and the Wellcome Trust (FC001194). This research was funded in whole, or in part, by the Wellcome Trust (grant FC001194). For the purpose of Open Access, the author has applied a CC-BY public copyright licence to any Author Accepted Manuscript version arising from this submission.

## Supplementary Data

**Figure S1.**
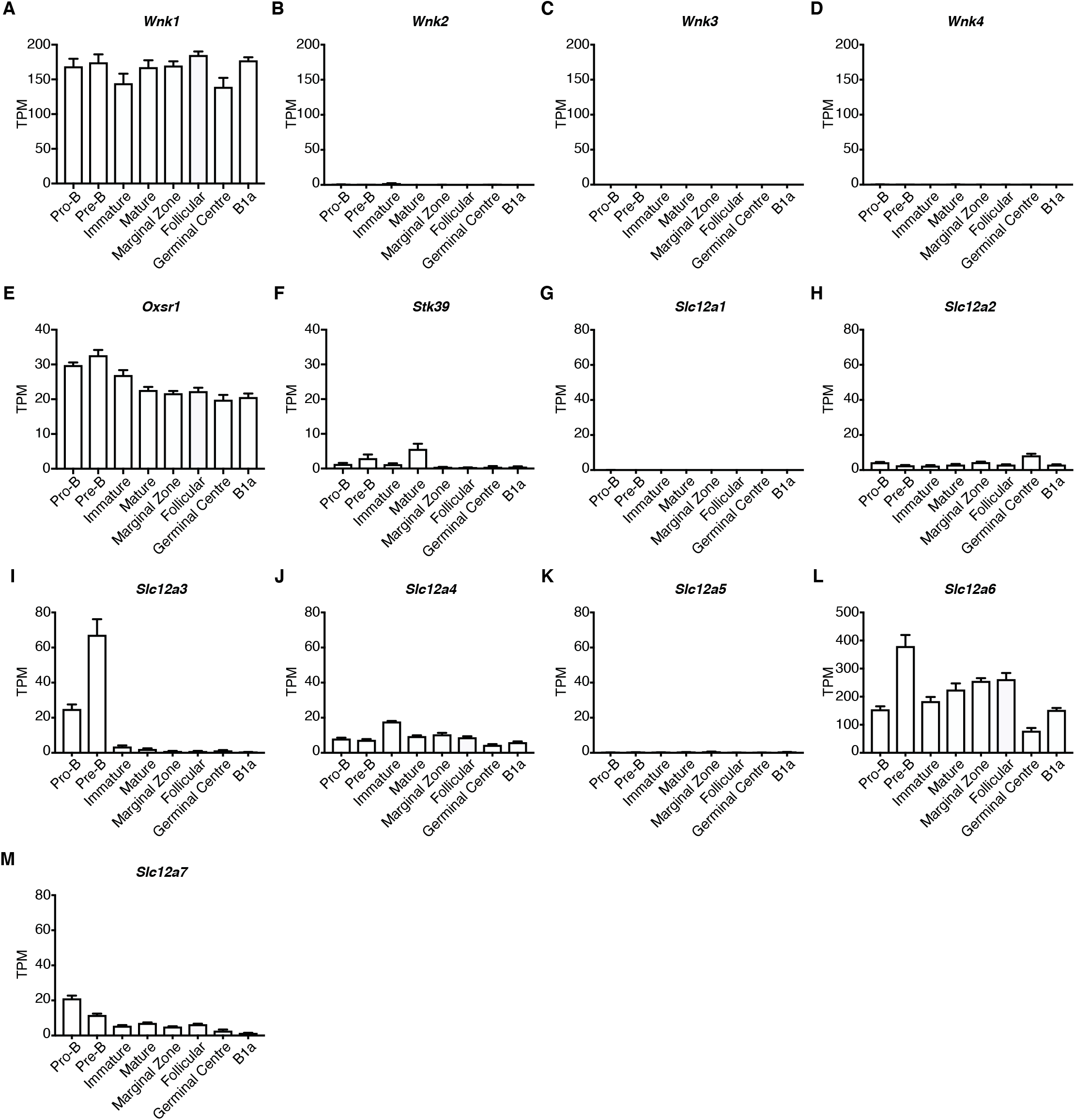
Expression of WNK pathway genes in B cell subsets. (A-M) Mean ±SEM expression levels of the indicated genes in pro-B, pre-B, immature B and mature B cells in bone marrow, marginal zone, follicular and germinal center B cells in spleen, and B1a cells from the peritoneal cavity as determined by RNAseq (Brazão et al., 2016). Expression is measured as transcripts per million (TPM). Sample size, n=5.

**Figure S2.**
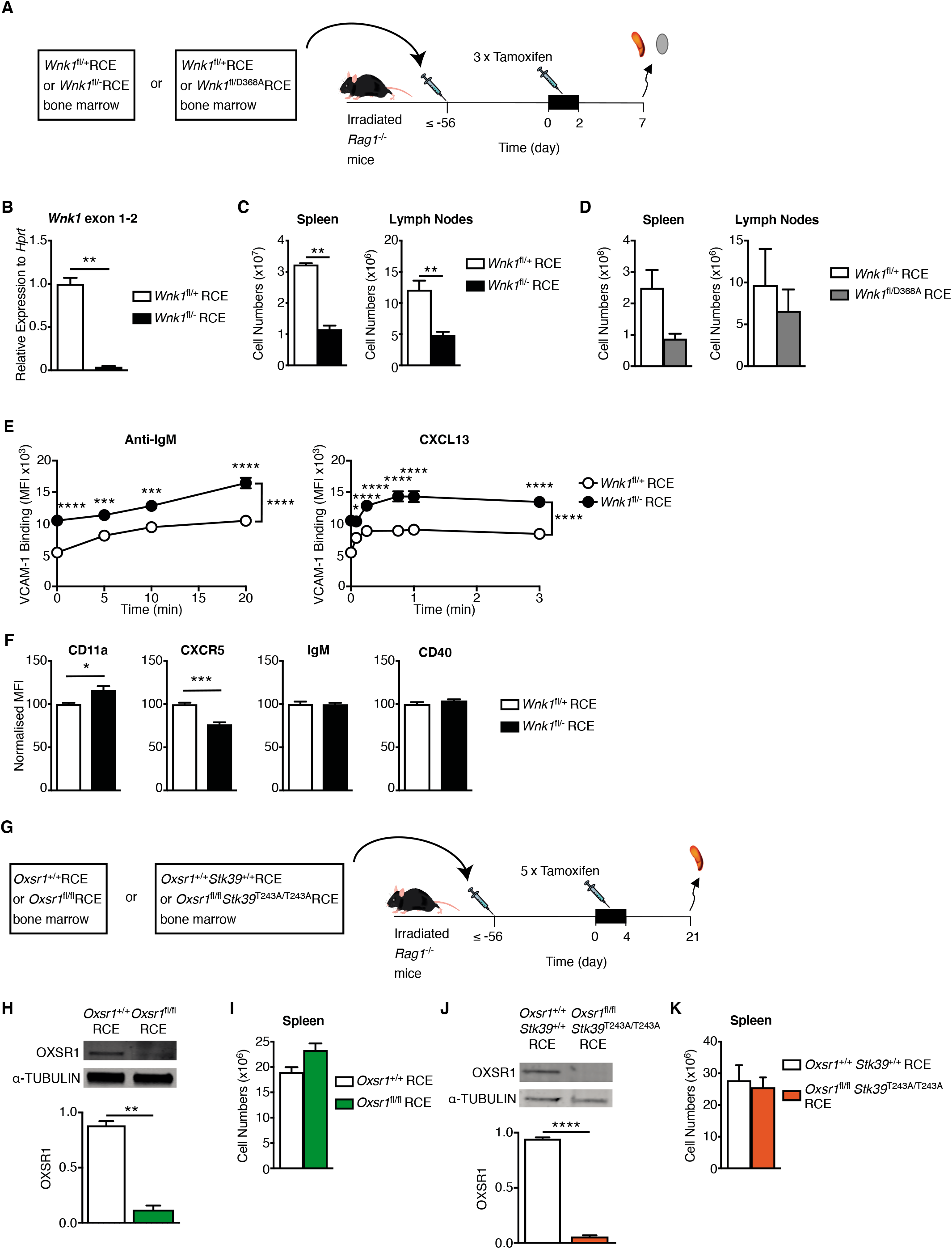
Characterization of mice with mutations in WNK1 pathway genes. (A) Irradiated RAG1-deficient mice were reconstituted with bone marrow from either *Wnk1*^fl/+^RCE or *Wnk1*^fl/fl^RCE mice, or from *Wnk1*^fl/+^RCE or *Wnk1*^fl/D368A^RCE mice. At least 56 d later, mice were treated with tamoxifen on 3 consecutive days and analyzed 7 d after start of tamoxifen treatment. (B) Mean±SEM levels of *Wnk1* mRNA measured across the junction of exons 1 and 2 in control or WNK1-deficient splenic mature B cells, normalized to *Hprt* expression and to *Wnk1* mRNA levels in control B cells (set to 1). (C, D) Mean±SEM numbers of mature B cells in the spleen (B220^+^CD19^+^CD93^-^) and lymph nodes (TCRβ^-^B220^+^IgM^+^IgD^+^) of RAG1-deficient mice reconstituted with *Wnk1*^fl/+^RCE or *Wnk1*^fl/fl^RCE marrow (C) or with *Wnk1*^fl/+^RCE or *Wnk1*^fl/D368A^RCE marrow (D), as described in A. (E) Mean±SEM binding of soluble VCAM-1 complexes to control or WNK1-deficient B cells in response to treatment with anti-IgM or CXCL13 for the indicated times. (F) Mean±SEM surface levels of CD11a, CXCR5, IgM and CD40 on control or WNK1-deficient B cells normalized to expression on control B cells (set to 100). (G) Irradiated RAG1-deficient mice were reconstituted with bone marrow from either *Oxsr1*^+/+^RCE or *Oxsr1*^fl/fl^RCE mice, or from *Oxsr1*^+/+^*Stk39*^+/+^RCE or *Oxsr1*^fl/fl^*Stk39*^T243A/T243A^RCE mice. At least 56 d later, mice were treated with tamoxifen on 5 consecutive days and analyzed 21 d after start of tamoxifen. (H, J) Immunoblot analysis (top) of total cell lysates from splenic B cells from RAG1-deficient mice reconstituted with *Oxsr1*^+/+^RCE or *Oxsr1*^fl/fl^RCE marrow (H) or with *Oxsr1*^+/+^*Stk39*^+/+^RCE or *Oxsr1*^fl/fl^*Stk39*^T243A/T243A^RCE marrow (J), probed with antibodies to OXSR1 and α-TUBULIN. Graph (bottom) shows mean±SEM amount of OXSR1 in the lanes above, normalized to the abundance of α-TUBULIN in each lane. (I, K) Mean±SEM numbers of mature B cells in the spleen (B220^+^CD19^+^CD93^-^) of RAG1-deficient mice reconstituted with *Oxsr1*^+/+^RCE or *Oxsr1*^fl/fl^RCE marrow (I), or with *Oxsr1*^+/+^*Stk39*^+/+^RCE or *Oxsr1*^fl/fl^*Stk39*^T243A/T243A^RCE marrow (K), as described in G. Mann-Whitney test (B, C, F, H-K); Two-way ANOVA (E), Mann-Whitney test (F, H-K); * 0.01 < *P* < 0.05, ** 0.001 < *P* < 0.01; *** 0.0001 < *P* < 0.001, **** *P* < 0.0001. Sample sizes: 6 (B, D, E), 5 (C, H), 7 (F, K), 3-4 (I), 12 (J). Data are pooled from 2 independent experiments.

**Figure S3.**
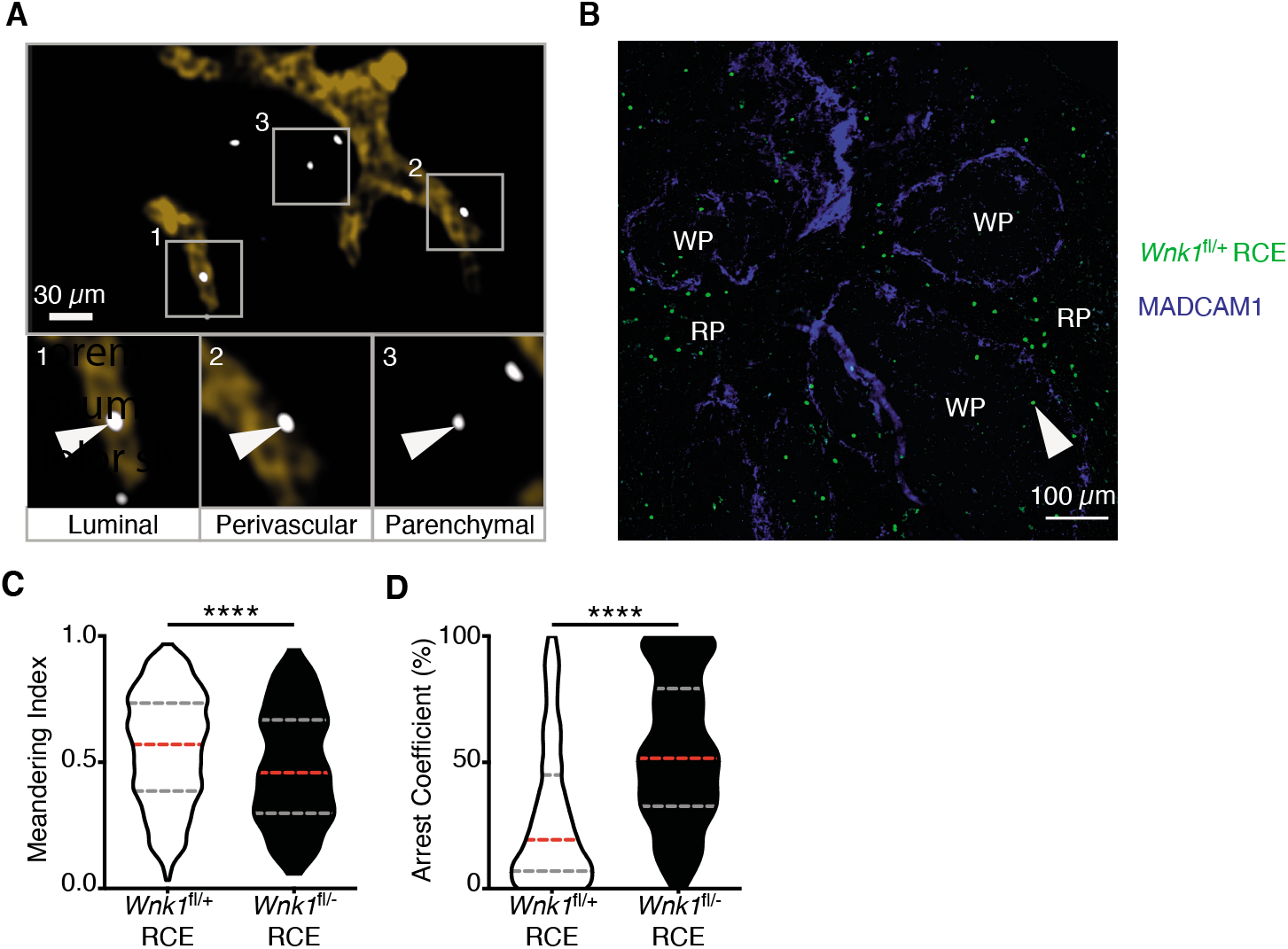
WNK1 is required for B cell homing and migration *in vivo*. (A) Example image from analysis of three-dimensional histology of lymph nodes 40 min after transfer of either wild-type or WNK1-deficient dye-labelled B cells into C57BL6/J mice, showing how the localization of transferred B cells was categorized as luminal, perivascular or parenchymal. White boxes (1-3) indicate areas that are enlarged below and white arrows indicate cells that fall into the indicated category. (B) Example image from analysis of histology of spleen 1 h after transfer of either control of WNK1-deficient B cells into C57BL/6J mice, showing transferred cells (green, indicated with white arrow) and MADCAM1 staining (blue). Cells in the white pulp (WP) were defined as cells within MADCAM1 staining, and cells in red pulp (RP) were defined as outside the MADCAM1 staining. (C, D) Analysis of migration of dye-labelled control and WNK1-deficient B cells in lymph node follicles by two-photon intravital microscopy 24 h after transfer into C57BL/6J mice. Violin plots of meandering index (C), a measure of track straightness, and arrest coefficient (D) defined as the proportion of time a cell moved <4 μm/min; dashed lines indicate median (red) and 25^th^ and 75^th^ percentiles (grey). Two-way ANOVA (A), Mann-Whitney test (B, D, E); **** *P* < 0.0001. Sample sizes: 5441 WNK1-deficient and 16196 control B cells (C, D). Data are pooled from 2 independent experiments.

**Figure S4.**
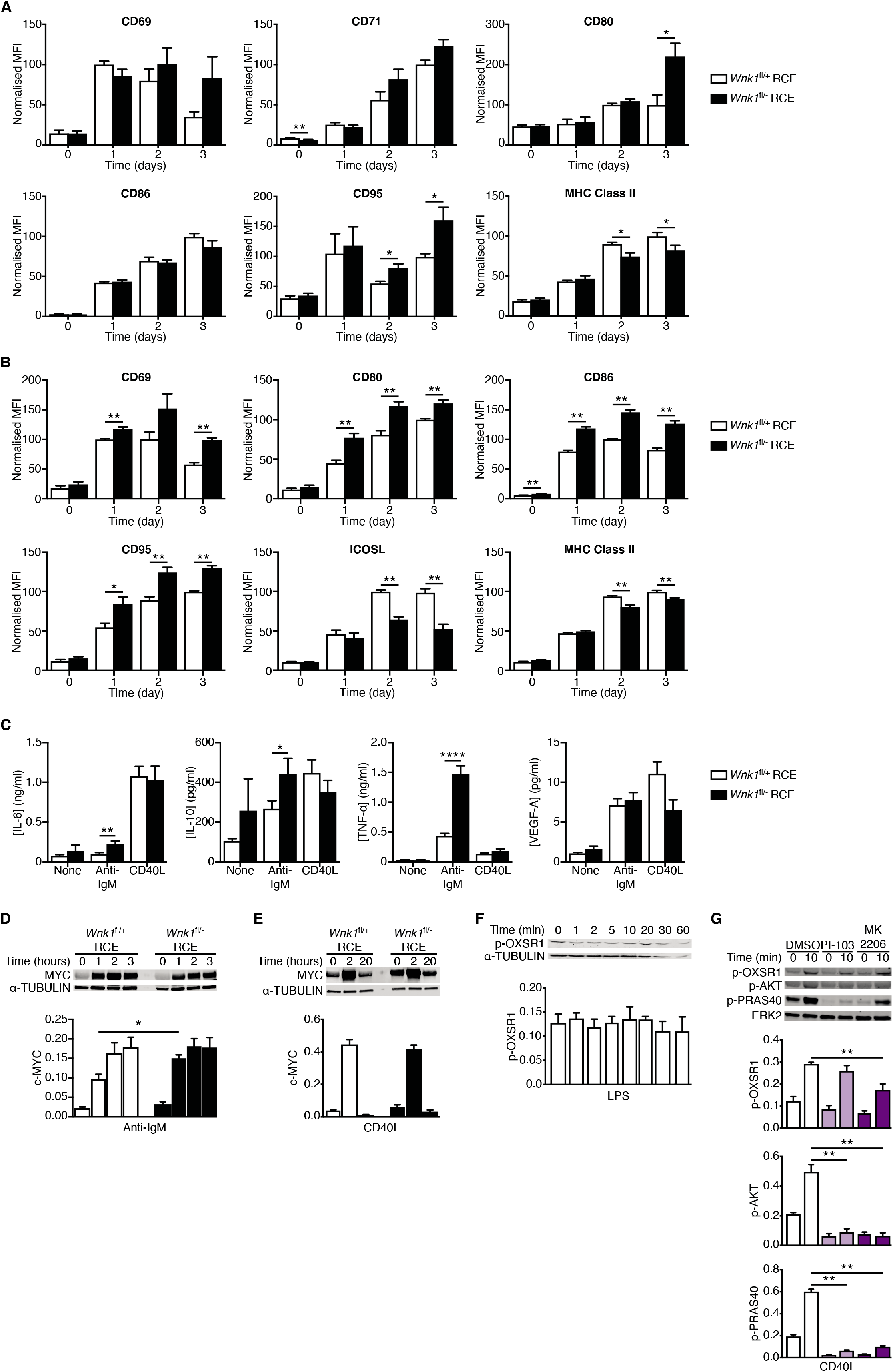
Loss of WNK1 results in altered levels of cell surface proteins on activated B cells. (A, B) Mean±SEM levels of the indicated cell surface proteins on control and WNK1-deficient B cells stimulated with anti-IgM (A) or CD40L (B) for the indicated times. Data shown as MFI normalized to the maximum response of control B cells (set to 100). (C) Mean±SEM concentrations of the indicated cytokines in medium from cultures of control or WNK1-deficient B cells that were unstimulated or stimulated with anti-IgM or CD40L for 72 h. (D and E) Immunoblot analysis (top) of cell lysates from control or WNK1-deficient B cells stimulated with anti-IgM (D) or CD40L (E) for the indicated times, probed with antibodies to MYC or α-TUBULIN. Graphs (below) show mean±SEM levels of MYC normalized to the abundance of α-TUBULIN in each lane. (F) Immunoblot analysis (top) of cell lysates from wild-type mouse B cells stimulated with LPS for the indicated times, probed with antibodies to p-OXSR1 and α-TUBULIN. Graph shows mean±SEM levels of p-OXSR1 normalized to the abundance of α-TUBULIN in each lane. (G) Top; immunoblots of total cell lysates from wild-type mouse B cells treated with vehicle (DMSO), a PI3K inhibitor (PI-103) or an AKT inhibitor (MK2206) and stimulated for the indicated times with CD40L, probed with antibodies to p-OXSR1, p-AKT, p-PRAS40 or ERK2. Below; graphs of mean±SEM abundance of p-OXSR1, p-AKT and p-PRAS40 in the lanes above, normalized to ERK2. Mann-Whitney test; * 0.01 < *P* < 0.05, ** 0.001 < *P* < 0.01, **** *P* < 0.0001. Sample sizes: 5-6 (A), 6 (B, D, E, G), 9-15 (C), 5 (F). Data are pooled from 2 (A, B, D, E) or 3 (C, F) independent experiments.

**Figure S5.**
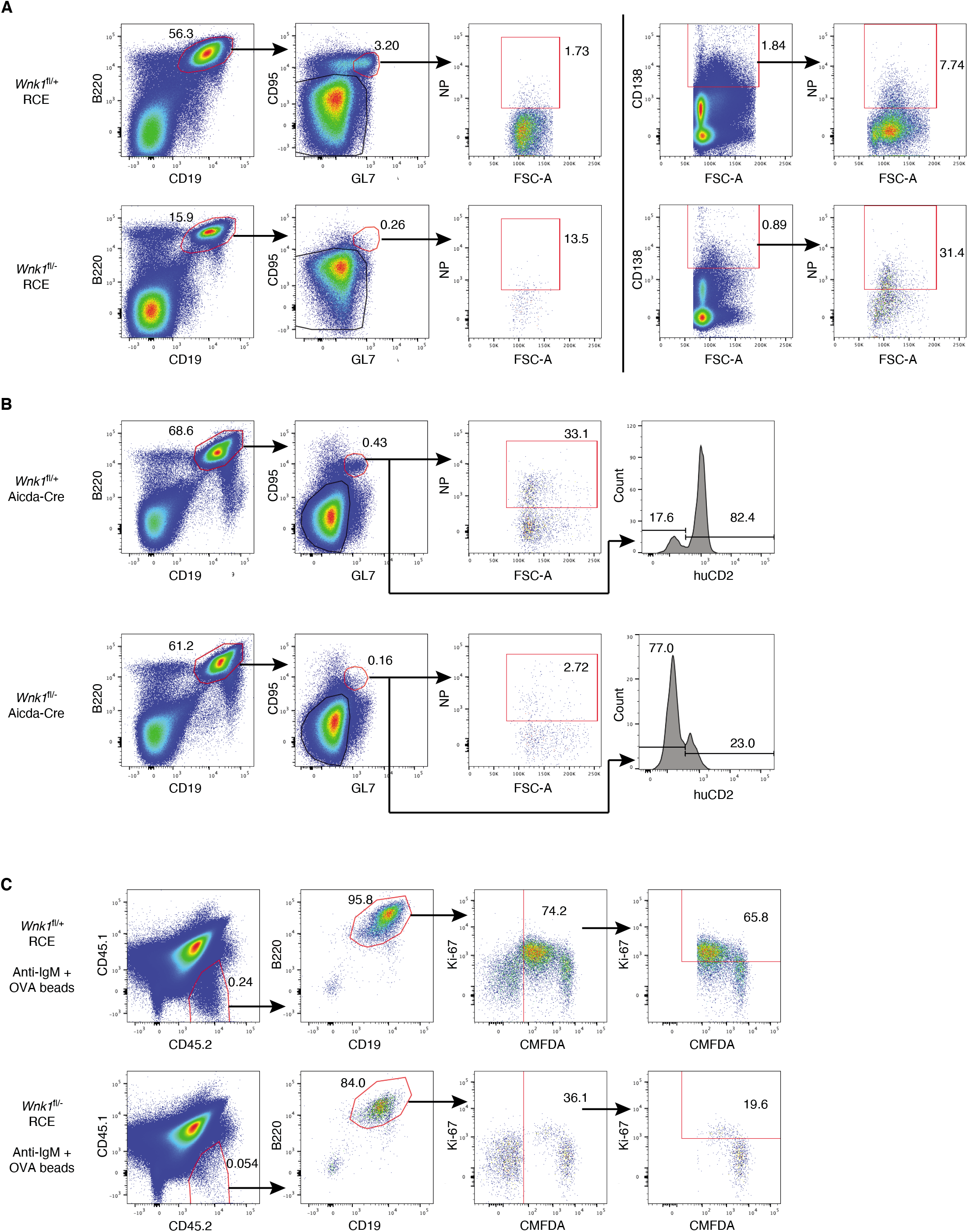
Loss of WNK1 results in failure to generate antigen-specific germinal center B cells and plasma cells and to to divide *in vivo*. (A) Flow cytometric analysis of splenocytes from mice immunized with NP-CGG in alum treated as described in Figure 6A, showing gating strategy for NP-specific germinal center cells (B220^+^CD19^+^CD95^+^GL7^+^NP^+^) and NP-specific plasma cells (CD138^hi^NP^+^) for control (top row) and mutant mice (bottom row). Numbers on dot plots indicate percentages of cell populations within gates (red boxes). (B) Flow cytometric analysis of splenocytes from mice immunized with NP-CGG in alum treated as described in Figure 6D, showing gating strategy for NP-specific germinal center cells (B220^+^CD19^+^CD95^+^GL7^+^NP^+^) and a histogram of human CD2 (huCD2) surface expression on germinal center B cells (B220^+^CD19^+^CD95^+^GL7^+^) for control (top row) and mutant mice (bottom row). Numbers in dot plots indicate percentages of cell populations within gates (red boxes), numbers on histogram indicate percentage of germinal center B cells that are negative or positive for huCD2 expression. (C) Flow cytometric analysis of splenocytes from mice treated as described in Figure 7A, showing gating strategy for control (top row) and mutant (bottom row) B cells that had been pre-treated with beads conjugated with both anti-IgM and OVA (CD45.1^-^ CD45.2^+^B220^+^CD19^+^CMFDA^+^) and transferred into a CD45.1^+^CD45.2^+^ host, showing analysis of % transferred B cells that were Ki67^+^. Numbers on dot plots indicate percentages of cell populations within gates (red boxes).

**Video S1**

Video showing representative examples of three-dimensional histology of lymph nodes 40 min after intravenous transfer of either wild-type (white cells, first half of video) or WNK1-deficient (blue cells, second half of video) B cells and 20 min after intravenous injection of MECA-79 antibody (orange). Analysis of B cell localization from these videos is shown in Figures 3C and S3A.

**Video S2**

A time-lapse video showing migration of B cells in lymph node follicles analyzed by two-photon intravital microscopy 24 h after transfer of control (white) and WNK1-deficient (blue) B cells into C57BL/6J mice from the experiment described in Figure 3E. Analysis of B cell tracks from these videos is shown in Figures 3F-3I and S3C and S3D.

**Video S3**

Video from two-photon intravital microscopy of the experiment described in Figure 7E, showing WNK1-expressing control B cells (blue), WNK1-deficient (KO) B cells (red) and GFP-expressing OT-II CD4^+^ T cells (green), with tracks showing migration of the B cells; transferred polyclonal CD4^+^ T cells are not shown. Time is shown in min:s. In the first part of the video the scale bar represents 20 μm, in the second higher magnification part of the video, the scale bar represents 10 μm.

**Video S4**

Video from two-photon intravital microscopy of the experiment described in Figure 7E, showing WNK1-expressing control B cells (blue), WNK1-deficient (KO) B cells (red) and GFP-expressing OT-II CD4^+^ T cells (green), showing interactions between OT-II T cells and control B cells (open arrowhead) or mutant B cells (filled arrowhead); transferred polyclonal CD4^+^ T cells are not shown. Time is shown in min:s. In the first part of the video the scale bar represents 20 μm, in the second higher magnification part of the video, the scale bar represents 10 μm.

